# Single-cell atlas of transcript usage remodelling in antiviral immune responses across human populations

**DOI:** 10.64898/2026.04.30.721889

**Authors:** Ruben Chazarra-Gil, Aida Ripoll-Cladellas, Maria Sopena-Rios, Iris Mestres-Pascual, Miquel Calvo, Ferran Reverter, Diego Garrido-Martín, Marta Melé

## Abstract

Humans exhibit substantial interindividual variation in their immune responses to infection, yet the contribution of transcript usage —the relative abundance of gene isoforms— to this variation remains poorly understood. Here, we generate the first single-cell atlas of transcript usage variation during early responses to influenza A virus and SARS-CoV-2 across 160 individuals of African and European ancestry. We show that viral stimulation induces widespread remodelling of transcript usage across all major immune lineages, with changes that are largely lineage-restricted and frequently undetected at the gene expression level. We further find that ancestry-associated effects on transcript usage are predominantly cell type-specific and contribute to population differences in antiviral responses. In addition, the genetic regulation of transcript usage during stimulation differs between influenza A and SARS-CoV-2, pointing to virus-dependent regulatory architectures. Together, our findings establish transcript usage as a dynamic regulatory layer shaping responses to viral infection across immune cell types and human populations, providing new insights into the molecular basis of variation in human antiviral immunity.

## Introduction

Transcript usage refers to the relative abundance of alternative isoforms expressed from the same gene. By modulating transcript usage, eukaryotic cells expand their transcriptome and proteome diversity through rearrangement of existing domains^1–3^. Multiple co- and post-transcriptional mechanisms shape transcript usage, such as alternative splicing, polyadenylation and transcription start site selection, generating transcript isoforms with distinct stability, regulatory elements, or protein domains^4–6^. As a result, transcript usage impacts virtually all cellular processes, from cell differentiation^7,8^ and development^9–11^ to tissue identity acquisition^1,12–14^ and disease^15–18^. In the immune system, transcript usage variation has been shown to fine-tune signal transduction and activation by generating receptor and adaptor isoforms with distinct signalling potential^19,20^. For instance, in macrophages, bacterial stimulation induces alternative transcript usage of the myeloid differentiation factor-2 gene, generating two isoforms with opposing effects: the long isoform enhances cytokine and type I interferon responses, while the short isoform dampens this response to prevent excessive inflammation^21,22^. Similarly, in T cells, antigen stimulation triggers a shift from longer to shorter CD45 isoforms that modulates T cell receptor signaling^23,24^. At transcriptome-wide scale, studies have shown extensive remodelling of transcript usage upon immune stimulation in monocytes^25^, macrophages^26–28^, and dendritic cells^29^ establishing transcript usage as a key feature of immune activation in response to pathogens.

Genetic variation is a major determinant of transcript usage differences between individuals^25,30–34^. Studies mapping splicing and transcript usage quantitative trait loci (sQTLs and TU-QTLs, respectively) have established the genetic basis of splicing variation^25,29,30,35–38^, showing that splicing-related QTLs are broadly shared across tissues^37,39^. At the single-cell level, a recent study showed that sQTLs often have consistent effect directions across immune cell types but different effect sizes, revealing substantial cell type– dependent genetic regulation of splicing under basal conditions^40^. In the context of infection, work in isolated myeloid cells has identified stimulation-specific genetic effects on splicing^25,28,41^, suggesting that some regulatory variants may become active only during pathogenic activation. However, the extent to which these infection-specific genetic effects generalize across immune cell types remains unclear. This underscores the need to map TU-QTLs in a cell type–resolved manner under different immune stimulations.

Single-cell transcriptomics has enabled systematic exploration of immune responses at cell type resolution across human populations, revealing cell type-specific gene expression programs that vary with genetic ancestries^42,43^. For instance, during influenza A virus (IAV) infection, donors of European ancestry show increased expression of interferon genes early after infection, which reduces viral activity at later stages^42^. Similarly, Aquino et al.^43^ found ancestry-related differences are largely driven by altered immune cell composition linked to latent cytomegalovirus infection. Together, these studies underscore the power of single-cell approaches to dissect ancestry-associated gene expression changes upon immune stimulation. In contrast, the impact of immune stimulation on transcript usage has been far less explored at single-cell resolution. A key limitation has been the difficulty of quantifying transcript usage from 3′ or 5′ end-biased single-cell RNA-sequencing (scRNA-seq) data, which generate short reads that are biased toward transcript ends and cannot be reliably assigned to isoforms. This limitation is particularly relevant because most population-scale single-cell cohorts rely on such end-biased technologies^40,42–46^. Consequently, how immune stimulation influences transcript usage across immune lineages and genetically diverse individuals remains largely unknown.

Here, we present the first comprehensive single-cell atlas of transcript usage variation in the immune response to IAV and SARS-CoV-2 (COV) infection across 160 genetically diverse individuals^43^. We first developed novel statistical frameworks for differential transcript usage (DTU) and TU-QTL analyses that robustly assign end-biased single-cell reads to transcript structures. Then, we applied these frameworks to identify infection-associated DTU across human populations and map TU-QTLs in five major immune cell lineages. Our results show that viral stimulation induces widespread remodelling of transcript usage across all immune lineages, and that such changes are highly lineage-specific and frequently independent from overall gene expression differences. We further find that ancestry-associated effects on transcript usage are predominantly cell type-specific and contribute to population differences in antiviral responses. Finally, we characterize genetic regulation of transcript usage across lineages in resting and stimulation contexts, and identify variants active only under a specific viral stimulation, suggesting pathogen-dependent genetic effects. Together, our study provides a cell type–resolved view of how viral infection reshapes isoform usage across immune cells in diverse human populations, and how genetic variation modulates this process, offering a framework to investigate isoform regulation during host–pathogen interactions.

## Results

### Equivalence class quantification captures viral-associated increase in transcriptome diversity

To systematically characterize transcript usage differences upon immune stimulation across the five major immune lineages (CD4^+^ and CD8^+^ T cells, monocytes, NK cells, and B cells), we analyzed a 3′ scRNA-seq dataset of 770,000 peripheral blood mononuclear cells (PBMCs) in baseline and stimulated with IAV and COV from 160 donors of European and African descent^43^ (Fig. 1a). Quantifying transcript usage from end-biased scRNAseq data (e.g., 3’ or 5’), however, presents a major technical challenge due to the lack of read coverage among most of the transcripts’ length (Fig. 1b). To overcome this limitation, we established a computational framework to quantify transcript usage in end-biased single cell data confidently. Specifically, we assigned all exonic reads to equivalence classes (ECs), which represent multi-sets of transcripts a read could have originated from^47–49^ (Fig. 1b). Importantly, because ECs represent aggregate transcript isoforms with read support, they provide a more robust quantification of transcript expression compared to probabilistic assignment of reads to specific transcripts^47–50^ when the uncertainty is extremely high. Consistent with this, transcript abundances inferred using the widely used expectation maximization (EM) algorithm^47–50^ showed poor agreement between bulk RNA-seq and donor-level pseudobulk scRNA-seq data, with significant correlations (false discovery rate (FDR)<0.05) observed for only 12% of the 12,136 transcripts detected in both datasets^42^ (mean ρ=0.13; <u>Methods</u>, <u>Supplementary Fig 1a</u>). Furthermore, gene-level visualizations revealed that reads mapping to multiple transcripts are often ambiguously assigned by the EM algorithm (Supplementary Fig 1b). Altogether, these results confirm that transcript abundance estimation is unreliable in 3’ scRNA-seq data, and support EC quantification as a more accurate alternative that preserves the underlying read-to-structure information based on coverage.

**Figure 1.**
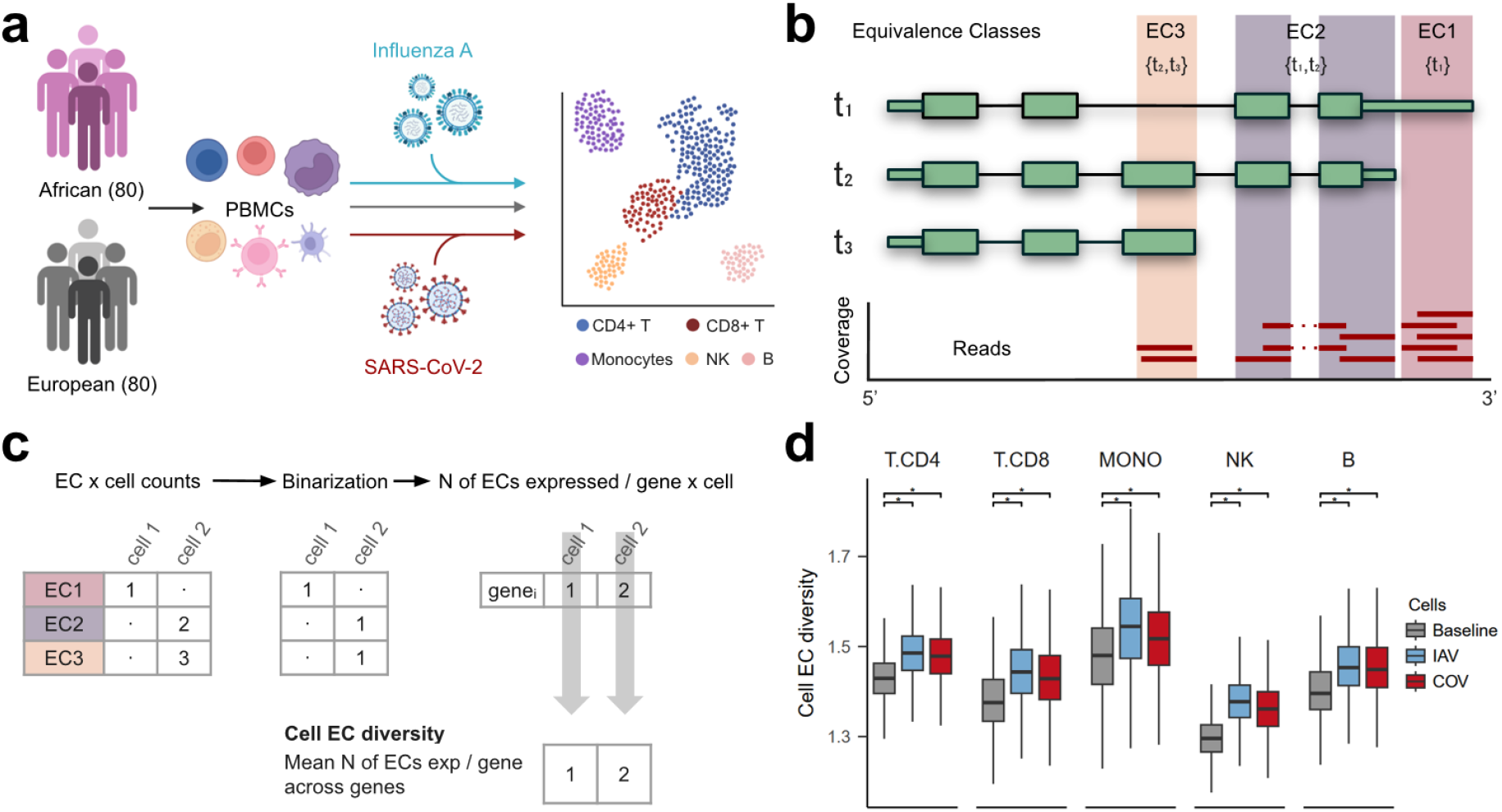
Viral stimulation increases transcriptome diversity across the five major immune lineages. **a)** 3’ Single-cell RNA sequencing (scRNA-seq) dataset of PBMCs (N=769,534) from 160 donors of European and African descent in baseline and stimulated with influenza A virus (IAV) and SARS-CoV-2 (COV)^43^. **b)** Schematic representation of equivalence class (EC) quantification in 3’ scRNA-seq data. Sequencing reads are assigned to ECs, each representing a segment of one or multiple transcripts. In the example gene shown, three ECs are generated: EC1 corresponds to the transcript 1-specific region in its 3’ end, while EC2 and EC3 correspond to regions shared between transcript 1 and 2 and transcript 2 and 3 respectively. **c)** Calculation of cell EC diversity metric to quantify viral-associated shifts in transcriptome complexity at single-cell resolution. **d)** Cell EC diversity is significantly higher in viral-stimulated (IAV,COV) compared to baseline cells (Wilcoxon test; FDR< 2.2 × 10^−16^).

Leveraging this EC-based framework, we next asked how immune response alters transcriptome complexity at single-cell resolution. Previous studies in monocytes and macrophages showed that immune stimulation increases transcriptome complexity, reflected in a higher diversity of expressed transcript isoforms per gene^25,26^. To investigate whether this occurs systematically across all immune lineages, we calculated, for each cell, the average number of expressed ECs (count > 0) per gene (‘cell EC diversity’) (<u>Methods</u>; Fig. 1c; <u>Supplementary Data 1a</u>). In all immune lineages, cell EC diversity was higher in viral-stimulated cells compared to baseline (Fig. 1d; FDR< 2.2 × 10^−16^). This increase was not explained by differences in sequencing depth between cells (<u>Methods</u>; <u>Supplementary Fig 2</u>, <u>Supplementary Data 1b</u>). Instead, the increase in transcriptome complexity in viral-stimulated cells was predominantly explained by highly upregulated genes (log_2_FC > 1) upon viral stimulation (<u>Methods</u>; <u>Supplementary Fig. 3</u>, <u>Supplementary Data 1c</u>), suggesting that transcriptional activation during viral stimulation is associated with increased isoform diversity.

Together, these results show that EC quantification in single-cell data robustly captures the infection-associated increase in transcriptome complexity across immune cell types, highlighting transcript diversity as a hallmark of the host response to viral challenge.

### Viral stimulation induces widespread transcript usage remodelling across immune cell types

Previous studies have shown that immune stimulation can alter transcript usage in myeloid cells^25–27,29^, but whether such remodelling occurs systematically across immune lineages remains less explored. To identify infection-associated transcript usage differences (infection DTU) across immune lineages, we developed a novel strategy based on multivariate non-parametric modelling of EC ratios^51^. Specifically, we subtracted the EC relative abundances between infected and baseline samples for each donor (e.g. Donor_1 IAV_-Donor_1Baseline_) and modelled the resulting EC difference vectors of a gene jointly (<u>Methods</u>; Fig. 2a). This approach allows to capture coordinated shifts in transcript usage while controlling for technical and biological covariates. Our framework yielded near-zero false positives when sample labels were randomly shuffled (mean=2.75 significant genes) (<u>Methods</u>; <u>Supplementary Fig. 4a</u>). We observed that viral stimulation induces extensive remodelling of transcript usage across all immune lineages (1,497 total unique infection DTU genes) (Fig. 2b; <u>Supplementary Data 2a</u>). The response was broader for IAV than for COV, with 47% more genes showing transcript usage shifts, consistent with the stronger transcriptional activation previously reported for IAV^43^. Monocytes had the largest number of infection-associated DTU genes, in line with a larger expression response to stimulation^42,43^, likely due to their recruitment and activation during viral infections^52–54^. To assess replication, we compared our monocyte infection DTU genes with an independent bulk monocyte splicing study under IAV infection^25^ and found significant overlaps (OR_IAV_=2.63, OR_COV_=2.99, FDR<4.38 × 10^−8^) (<u>Methods</u>; <u>Supplementary Fig. 4b</u>).

**Figure 2.**
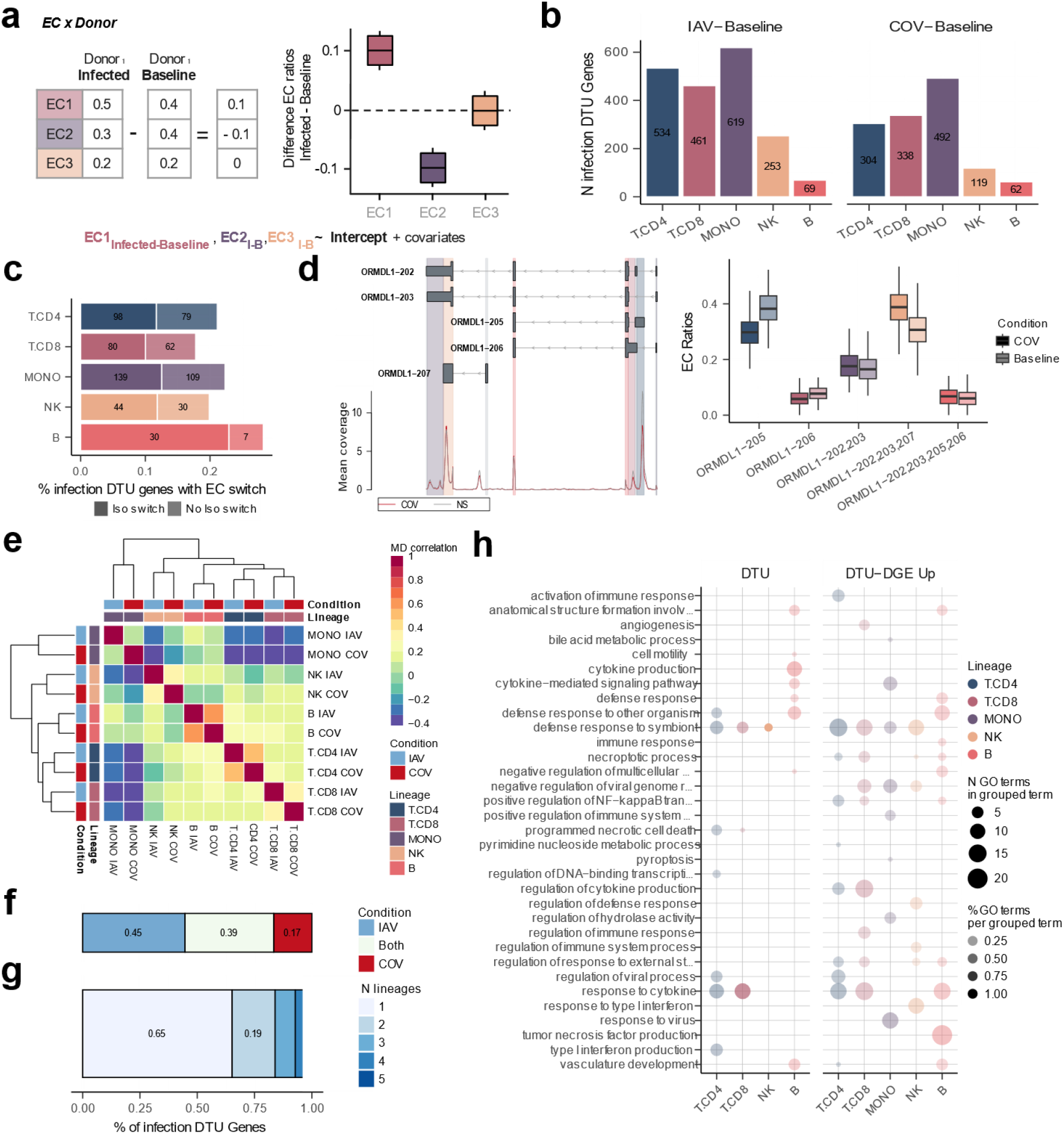
Viral stimulation triggers lineage- and virus-specific transcript usage remodelling across immune cell types. **a)** Infection differential transcript usage framework overview. For each gene within each immune lineage, we calculated donor-matched equivalence class (EC) differences between infected and baseline samples (e.g. Donor_1 infection_-Donor_1 Baseline_), and modelled the resulting EC difference vector jointly while accounting for technical and biological covariates. Genes with a significant intercept term (FDR < 0.05) are considered infection DTU genes. **b)** Number of infection DTU genes upon influenza A virus (IAV) and SARS-CoV-2 (COV) stimulation across immune lineages. **c)** Proportion of infection DTU genes with a switch in their mean most abundant EC between infected and baseline conditions. Most EC switches correspond to isoform switches (Iso switch), defined as changes between ECs mapping to non-overlapping transcript sets. **d)** Example of an isoform switch in an infection DTU gene. Left: Mapping of *ORMDL1* ECs to transcript regions. The five ECs correspond to distinct transcript fragments from five isoforms, represented in grey with UTRs shown as thin lines and coding sequences as thick lines. Colored vertical blocks highlight the transcript segments represented by each EC, matching the colors of the boxplots on the left. Regions colored in light grey represent lowly expressed ECs that were filtered. At the bottom, mean coverage profiles across COV and baseline samples are shown per genomic position (10-bp bins). Right: Relative abundances of *ORMDL1* ECs in B cells upon COV infection, showing a shift from ORMDL1-205 at baseline to ORMDL1-202/203/207 upon infection. **e)** Heatmap of infection DTU gene sharing based on the spearman correlation of maximum absolute difference (MD value) considering genes significant in at least one of the lineage-response combinations. Hierarchical clustering of lineage-response pairs is displayed. **f)** Sharing of infection DTU genes across IAV and COV responses considering genes tested in both responses for each immune lineage separately. **g)** Sharing of infection DTU genes across immune lineages considering genes tested in all lineages for each response separately. **h)** Pathway enrichment analysis of COV response infection DTU genes. Two gene sets are considered: i) all infection DTU genes, ii) DTU genes shared with upregulated infection differentially expressed genes and use all genes tested in the DTU analysis as background (Hypergeometric test). GO biological terms (FDR<0.05) were grouped using rrvgo^58^. Dot shade intensity and size reflects the fraction and absolute number of GO terms per grouped terms, respectively.

Because immune activation is known to induce isoform switching^21–24^, we examined whether infection DTU genes showed changes in the dominant isoform. Following infection, 16–29% of DTU genes shift to an alternative dominant EC. Over half of these events (57.5%) correspond to switches in the most abundant isoform (Fig. 2c), as the switching ECs map to distinct transcript sets —for example, the *ORMDL1* gene in B cells upon COV infection (Fig. 2d). Our findings reveal that isoform switching is pervasive across immune lineages during viral infection, consistent with prior evidence that this mechanism is involved in immune response modulation^19,20,55–57^.

Next, we examined whether transcript usage responses to infection were shared between viral responses across different immune lineages. To do this, we quantified the magnitude of each response as the maximum absolute difference (MD value) in mean EC ratio across baseline and stimulated conditions, and correlated responses across lineages (<u>Methods</u>). Overall, transcript usage responses exhibited moderate correlation between the two viral responses within each lineage (mean Spearman *ρ*=0.36) (Fig. 2e; <u>Supplementary Fig. 5</u>). Specifically, only 39% of infection DTU genes were shared across viruses, while 45% and 11% were specific to IAV and COV, respectively (Fig. 2f). Monocytes showed a substantially greater heterogeneity (mean *ρ*=0.05), consistent with previously reported virus-specific transcriptional responses in myeloid cells^43^. The moderate overlap between the two viral responses at the level of transcript usage contrasts with gene expression responses, which were far more shared across viruses, with 62% of differentially expressed genes (DEGs) common to both infections (<u>Supplementary Fig. 6a</u>; <u>Supplementary Data 2b</u>).

We then evaluated lineage specificity upon stimulation and found that most infection DTU genes (65%) were detected in a single immune lineage, indicating strong lineage restriction (Fig. 2g; <u>Supplementary Fig. 6b</u>). Importantly, this pattern persisted not only in higher-powered lineages (such as CD4+ and CD8+ T cells and monocytes) but also in smaller cell types (NK and B cells), where many infection DTU genes remained lineage-specific (<u>Supplementary Fig. 6c</u>). In contrast, only 45% of infection DEGs were lineage-specific (<u>Supplementary Fig. 6b,d</u>). Consistent with this, infection DEGs (|log_2_FC| > 0.5, FDR < 0.01) were significantly more likely to also exhibit DTU (Fisher’s exact test; Odds Ratio (OR) > 3.3, FDR < 1.05 × 10^−16^), yet a substantial fraction of infection DTU genes (29–59%) did not change in overall expression upon infection (<u>Supplementary Fig. 8</u>). Together, these findings indicate that, despite broad overlap in gene expression responses across cell types and viruses, transcript usage is more variable and often remodeled in a lineage- and response-specific manner. While part of this specificity may reflect the lower statistical power of DTU compared with differential gene expression analyses, the observation that the overall pattern holds in lower-powered cell types, emphasizes the importance of DTU in capturing cell type regulatory differences not evident at the gene expression level.

Finally, to assess the functional impact of these transcript-level changes, we investigated the biological pathways associated with infection DTU genes. We observed a generalized enrichment in immune response processes, including cytokine and interferon-related pathways (Fig. 2h; <u>Supplementary Fig. 8,9</u>, <u>Supplementary Data 2c-e</u>). Notably, we observed a larger number of enrichments for DTU genes that were also upregulated at the gene expression level (Fig. 2h), indicating that coordinated changes in expression and transcript usage preferentially affect core antiviral programs across immune lineages. Thus, although DTU events are often lineage-specific, their functional consequences converge on shared antiviral pathways, highlighting transcript usage as an additional layer of regulation that modulates conserved immune processes during viral infection.

### Genetic regulation is a major determinant of ancestry-associated transcript usage differences

Genetic ancestry is a main driver of transcript usage variation^25,30–34,38^. Yet, the study of transcript usage differences between individuals of genetically diverse populations across different stimulation conditions has remained limited to isolated cell types^25,36^. To identify population-specific transcript usage differences (population DTU) across immune cell types, we compared EC ratios between individuals of African and European ancestries within each immune lineage and condition (e.g. CD4+T-Baseline, CD4+T-IAV, CD4+T-COV), while controlling for technical and biological covariates^51^ (<u>Methods</u>; Fig 3a). Ancestry-associated differences in transcript usage were widespread across immune lineages and conditions, with a total of 438 unique population DTU genes identified after correcting for interindividual differences in cell composition^43^ (Fig. 3b; <u>Supplementary Data 3a</u>). Notably, adjusting for cell abundances reduced population DTU genes by 22.5–89.5%, particularly in NK cells and monocytes (mean reduction across conditions: 89.5% and 60.7%, respectively), consistent with pronounced ancestry-associated differences in subcellular composition within these lineages^43^. Most ancestry-associated DTU effects were highly lineage-specific, with 57.5–70.5% detected in a single immune lineage across baseline or stimulation conditions (Fig 3c; <u>Supplementary Fig. 10a</u>). In addition, a large fraction of population DTU genes (38.5-55.6%) were identified only after viral stimulation (<u>Supplementary Fig. 10a,b</u>). Consistently, ancestry-associated gene expression changes exhibited strong lineage specificity (70-75%), with a comparable fraction of population-associated DEGs emerging only upon infection (32.9–64.6%) (<u>Supplementary Fig. 10c-e</u>).

**Figure 3.**
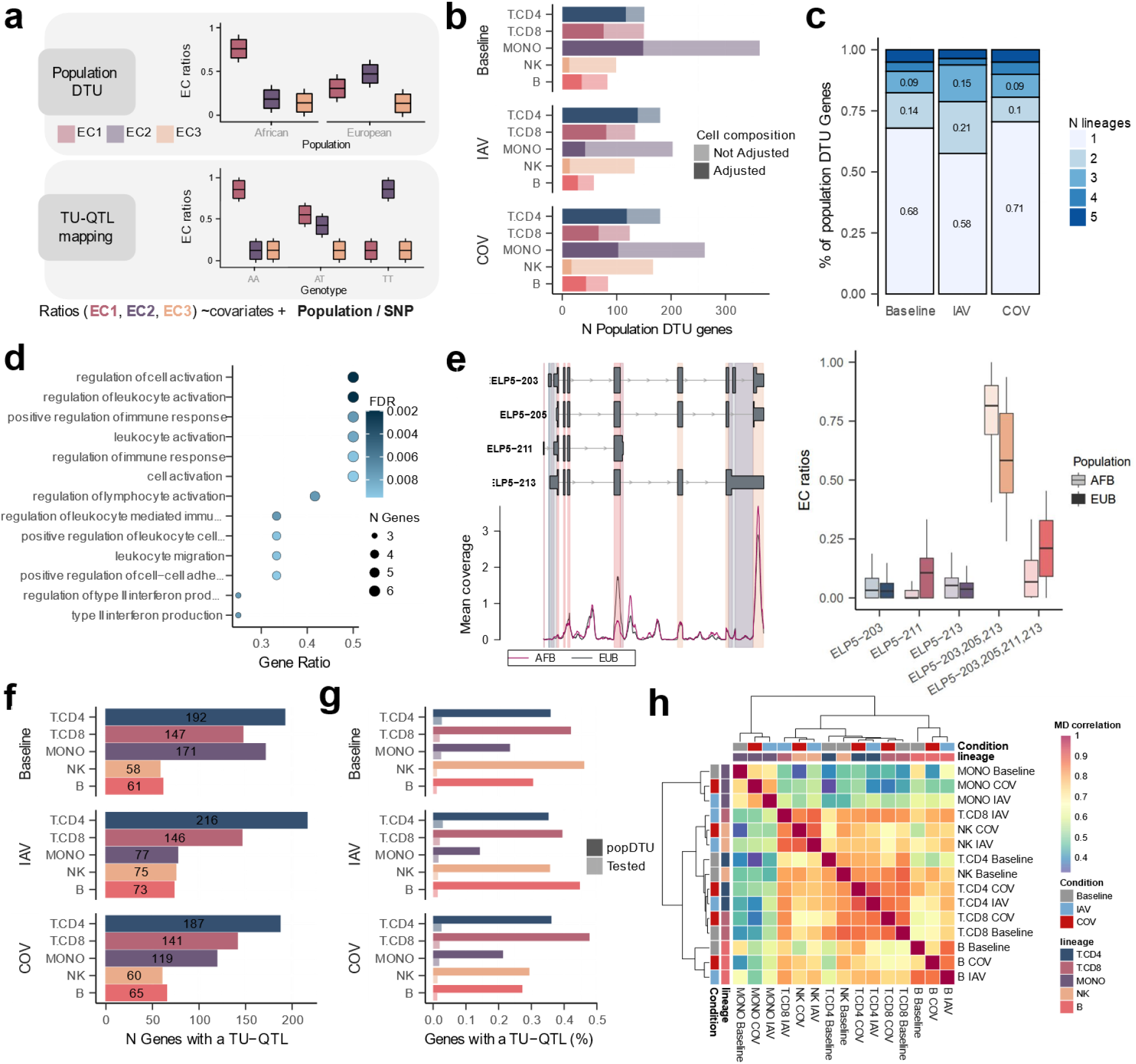
Ancestry-associated differences in transcript usage are lineage-specific and under genetic control. **a)** Schematic of the population differential transcript usage (DTU) and transcript usage quantitative trait loci (TU-QTL) analyses. For each gene, within each immune lineage and condition, EC ratios were modeled jointly across groups (e.g. populations or genotypes) while accounting for technical and biological covariates. **b)** Number of population DTU genes across lineages and conditions detected (FDR<0.05) with and without adjusting for cell composition. **c)** Sharing of population DTU genes across immune lineages, considering only genes tested in all lineages within each condition. **d)** Pathway enrichment analysis for baseline NK cell population DTU genes (N=13; Hypergeometric test, FDR < 0.01). Gene Ratio indicates the fraction of DTU genes present in the enriched term gene set. **e)** Example of a population DTU gene (*ELP5*) in baseline NK cells enriched for cell migration (GO:0016477) and cell motility (GO:0048870). Left: Mapping of *ELP5* ECs to transcript regions. The five ECs correspond to distinct transcript fragments from four isoforms, represented in grey with UTRs shown as thin lines and coding sequences as thick lines. Colored vertical blocks highlight the transcript segments represented by each EC, matching the colors of the boxplots on the right. Grey vertical blocks represent ECs filtered before the DTU analysis. At the bottom, mean coverage profiles across European and African individual samples are shown per genomic position (10-bp bins). Right: Relative abundances of the *ELP5* ECs by population. **f)** Number of genes with a TU-QTL across lineages and conditions. **g)** For each immune lineage and condition, the bars indicate the percentage of genes with a significant TU-QTL among population DTU tested and significant genes, respectively. **h)** Heatmap of TU-QTLs sharing based on the spearman correlation of maximum absolute difference (MD value) considering gene-SNP pairs significant in at least one of the lineage-condition comparisons. In addition, hierarchical clustering of lineage-condition pairs is displayed.

Because changes in dominant transcript isoforms can have functional consequences^21–24,59,60^, we asked whether ancestry-associated DTU was accompanied by these shifts. Notably, 12.9% of ancestry-associated DTU events involved a switch in their most abundant EC (<u>Supplementary Fig. 11</u>), reflecting transcript changes that may alter protein domains or regulatory regions. One example is the immunoglobulin A (*IgA*) heavy-chain gene (<u>Supplementary Fig. 12</u>), where population differences in transcript usage shift the balance between membrane-bound and secreted *IgA* proteins^61–63^ in B cells, consistent with known population diversity in immunoglobulin heavy-chain variable regions^64,65^.

To assess the functional relevance of the identified ancestry-associated transcript usage differences, we examined whether population DTU genes were enriched for specific biological processes. Population-DTU genes in NK cells at baseline showed particularly strong enrichment for leukocyte activation (FDR<1.9×10^−3^), regulation of type II interferon production (FDR<7.3×10^−3^) and leukocyte migration (FDR<9.2×10^−3^) (Fig. 3d; <u>Supplementary Fig. 13</u>, <u>Supplementary Data 3c</u>) such as the Elongator complex subunit gene *ELP5* —a regulator of cell migration and motility (GO:0016477; GO:0048870)— that showed population-specific transcript usage in NK cells (Fig. 3f). To confirm robustness of our framework, donor labels were permuted, yielding near-zero significant population DTU genes (mean=1.64) (Methods; <u>Supplementary Fig. 14a</u>). We further validated monocyte population DTU genes using an independent bulk monocyte dataset (Methods; <u>Supplementary Fig. 14b</u>).

Next, to investigate the extent to which population differences in transcript usage are driven by population differences in allele frequencies, we first mapped transcript usage quantitative trait loci (TU-QTLs) —jointly in both populations— by testing genetic variants for association with EC ratio shifts across immune lineages and conditions (Fig 3a; <u>Methods</u>). At a 1% FDR, we identified 1,385-6,698 TU-QTLs, affecting 58-216 genes (1.3-3.9% of the tested genes; Fig 3g; <u>Supplementary Fig. 15</u>, <u>Supplementary Data 3b</u>). Notably, population DTU genes were substantially more likely to harbor at least one TU-QTL at both baseline and stimulated conditions (Fig 3g; Fisher’s exact test, OR>10.7, FDR< 5.42×10^−5^). These associations significantly overlapped with those detected across multiple independent datasets, including a 5’ scRNA-seq PBMC cohort from Asian donors^40^ (Fisher’s exact test; OR>2.9, FDR<0.02) and three bulk RNA-seq datasets spanning monocytes^25^ (Fisher’s exact test; OR>5.47, FDR<1.46×10^−12^), lymphocytes (Fisher’s exact test; OR>5.67, FDR<3.85×10^−12^) and whole blood^37^ (Fisher’s exact test; OR>4.97, FDR<2.25×10^−13^) (<u>Supplementary Fig. 16</u>). Furthermore, population DTU genes with the largest ancestry differences were more likely to be under genetic regulation and associated with large-effect TU-QTLs (<u>Supplementary Fig. 17</u>), in line with previous observations at the gene expression level^43^. Next, we investigated whether genetic regulation influences the degree of lineage and condition specificity observed for ancestry-associated TU differences. Population DTU genes that harbored a TU-QTL showed substantially greater sharing across biological contexts (immune lineages and across stimulation conditions) than genes lacking TU-QTLs (<u>Supplementary Fig. 18</u>). Furthermore, clustering of TU-QTL effect sizes revealed a strong concordance of genetic effects across lineages and contexts (mean ρ=0.671) (Fig 3h). This concordance increased further after excluding monocytes (mean ρ=0.761), consistent with their weaker correlation with other lineages (mean ρ=0.5) as a result of higher transcriptional heterogeneity (Fig. 2e). Consistent with this high degree of genetic sharing, 13-17% of TU-QTLs identified within each condition were shared across all immune lineages, and 31–64% of TU-QTL variants were common to both baseline and stimulated conditions (<u>Supplementary Fig. 19</u>). Overall, these findings suggest that genetic regulation contributes primarily to ancestry-associated transcript usage differences that are shared across contexts, whereas more specific effects may be driven by non-genetic factors.

### Viral stimulation reveals context-dependent genetic effects on transcript usage

Gene expression responses to viral stimulation differ across populations due to both environmental and genetic factors^36,42,43,66^. However, differences between populations in transcript usage response to stimulation remain largely unexplored. To address this, we evaluated population × infection interaction effects by modelling donor-matched differences in EC ratio between infected and baseline samples (e.g. Donor_1 IAV_-Donor_1Baseline_), incorporating a population term in the model and adjusting for cell composition differences between individuals (<u>Methods</u>). We detected population-specific differences in transcript usage response to stimulation across all immune lineages, with the strongest signal observed in monocytes responding to COV stimulation (Fig.4a; <u>Supplementary Data 4a</u>). This pattern is consistent with previously reported ancestry-associated differences in monocyte gene expression response to COV^43^, likely reflecting the prominent monocyte response to pathogenic stimulation^42,43,52–54^. Despite limited statistical power, we also identified population-dependent transcript usage responses in other immune lineages. For instance, *KLRC1* —an inhibitory NK receptor that regulates CD8+ T cell exhaustion during viral infection^67,68^ — exhibited population-dependent shifts in isoform usage (*KLRC1-201/202* vs. *KLRC1-206*) upon IAV infection in CD8+ T cells (Fig.4b). We confirmed the robustness of our framework by permuting donor labels, yielding near-zero significant population DTU genes (mean=0.53; <u>Methods</u>; <u>Supplementary Fig. 20</u>).

Given the context-dependent genetic architecture of alternative splicing^25,28,41^, we sought to identify genetic effects on transcript usage that modify immune responses to viral stimulation (response TU-QTLs, reTU-QTLs). Restricting our analysis to genes expressed in both resting and stimulation conditions and to variants significantly associated with transcript usage only after infection (FDR_IAV,COV_ < 0.01, p_Baseline_ > 0.01) (Methods), we identified 1,279 reTU-QTLs in 108 genes across immune lineages (Fig 4c; <u>Supplementary Data 4b</u>). Most reTU-QTL were detected in a single cell type (~88%; Supplementary Fig. 21), indicating that stimulation-dependent genetic effects on transcript usage are likely restricted to one lineage. Furthermore, reTU-QTLs were also frequently virus-specific (Fig 4c), with monocytes showing the strongest virus-dependent signal (95.4%), in line with previous observations of pathogen-dependent splicing regulation in myeloid cells^25^. For example, upon IAV infection, the strongest response-specific reTU-QTL (rs592261, MD value_IAV-Baseline_=0.1, FDR_IAV_<4.3×10^−3^, MD_COV-Baseline_=0.03, p_COV_>0.55, p_Baseline_>0.94) (Fig 4d,e) was identified in monocytes at *ELP2* (Elongator Complex Protein 2), a gene involved in transcriptional elongation and histone modifications^69^. As IAV hijacks the host transcriptional machinery and requires transcription elongation for replication^70^, variants that alter *ELP2* splicing specifically during IAV infection may modulate host transcriptional capacity in a virus-specific manner. In contrast, upon COV infection, the strongest response-specific signal (rs116567297, MD_COV-Baseline_=0.11, FDR_cov_<3×10^−4^, MD_IAV-Baseline_=-0.01, p_IAV_>0.63, p_Baseline_>0.37) (Fig 4d,f) was detected in CD8+T cells at *NOTCH2* (Notch Receptor 2), a regulator of CD8+T cell differentiation^71^. NOTCH signalling, a key pathway in COVID-19 pathogenesis^72–74^, critically determines whether CD8+ T cells adopt effector versus regulatory phenotypes. Thus, genetic variants altering *NOTCH2* splicing specifically during COV infection may thereby influence this balance. Overall, these results indicate that viral infection reveals lineage- and pathogen-dependent genotype-by-environment effects on transcript usage.

**Figure 4.**
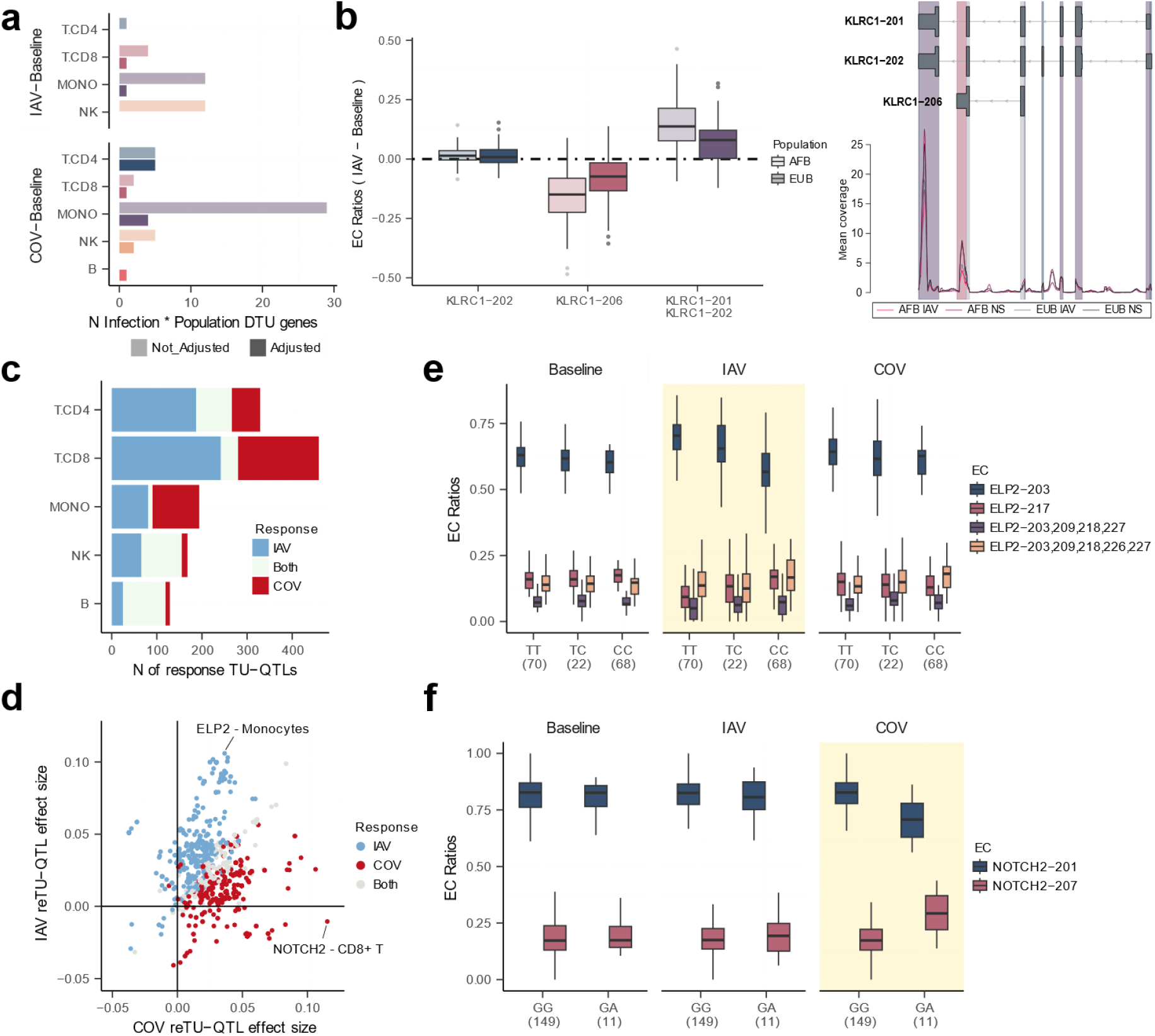
Genetic regulation of transcript usage response to stimulation is virus-specific. **a)** For each immune lineage and viral response, number of significant population × infection transcript usage interactions detected (FDR<0.1), shown with and without adjustment for interindividual differences in cell composition. **b)** Example of a population × infection transcript usage interaction in *KLRC1* in CD8+ T cells upon influenza A virus (IAV) infection. Left: Donor-matched differences in *KLRC1* equivalence class (EC) ratios between IAV and baseline samples, stratified by population. Right: Mapping of *KLRC1* ECs to transcript regions. The three ECs correspond to distinct transcript fragments from three isoforms, represented in grey with UTRs shown as thin lines and coding sequences as thick lines. Colored vertical blocks highlight the transcript segments represented by each EC, matching the colors of the boxplots on the left. Grey vertical blocks represent ECs filtered before the differential transcript usage analysis. At the bottom, mean coverage profiles across IAV and baseline samples in European and African individuals are shown per genomic position (10-bp bins). **c)** For each immune lineage, number of response transcript usage quantitative trait loci (reTU-QTLs), defined as SNPs that are significant TU-QTLs in infected conditions (FDR<0.01) but not at baseline (p>0.01). **d)** Comparison of reTU-QTL effect sizes between SARS-CoV-2 (COV) (MD_COV-NS_) and IAV (MD_IAV-NS_). Each point represents a single reTU-QTL (SNP–gene–lineage combination), colored by the viral condition in which the effect is detected. **e)** Example of an IAV-specific reTU-QTL at *ELP2* (rs1010955) in monocytes. **f)** Example of a COV-specific reTU-QTL at *NOTCH2* (rs116567297) in CD8+ T cells.For boxplots in panels **b, e** and **f**, center lines indicate medians, notches the 95% confidence intervals of the median, boxes the interquartile range, and whiskers 1.5× the interquartile range.

Finally, we asked whether these stimulation-specific genetic effects could contribute to population differences in transcript usage responses to infection. To this end, we evaluated whether genes harbouring reTU-QTLs overlapped with those with significant population *x* infection interaction effects. Direct gene-level overlap was limited, likely due to low statistical power in the population x infection interaction DTU analysis (Fig. 4a). However, population DTU genes were significantly enriched for reTU-QTL under stimulated conditions compared to baseline (Fisher’s exact test; OR_IAV,COV_>9.17, FDR_IAV,COV_<0.01, OR_Baseline_>0, FDR_Baseline_>0.77, <u>Supplementary Fig. 22</u>), suggesting that loci displaying population differences in transcript usage are more likely to harbor stimulation-specific genetic variants.

Together, these results reveal that viral stimulation uncovers context-dependent genetic regulation of transcript usage that may partially contribute to population diversity in antiviral immune response across immune cell types.

## Discussion

In this study, we present the first comprehensive single-cell atlas of transcript usage variation during early response to viral infection in immune cells across human populations. To address this, we developed novel statistical frameworks for DTU and TU-QTL analyses tailored to end-biased scRNA-seq data. As most large single-cell cohorts are 3’ or 5’ end-biased^40,42–46,75^, isoform quantification remains challenging. Our approach leverages EC quantification to robustly capture transcript usage variation in end-biased single-cell datasets, which enables the systematic investigation of isoform regulation across large-population cohorts and diverse biological contexts.

Using these frameworks, we find that viral infection induces widespread transcript usage remodelling across all major immune lineages, consistent with previous reports of global splicing shifts following bacterial and viral stimulation in myeloid cells^25–29^. Most infection-associated DTU genes are predominantly lineage-specific. Although differences in statistical power may contribute to this pattern, the detection of lineage-specific DTU genes in minor cell types suggests that lineage specificity reflects true biological signals. Moreover, a large fraction of transcript usage immune responses are virus-specific, indicating that IAV or COV differentially modulate isoform usage programs. This pattern contrasts with the broader sharing observed for gene expression differences upon viral stimulation across immune lineages, where antiviral transcriptional responses are more similar across lineages and stimulations^42,43^. Together, these findings suggest that transcript usage provides a more context-dependent regulatory layer that enables fine-tuning of cellular function while maintaining conserved antiviral gene expression programs^42,43^.

Consistent with these findings, we also observed limited overlap between differential transcript usage and gene-level expression changes. Many infection-associated DTU events occurred without corresponding changes in overall gene abundance, indicating that transcript usage remodelling captures aspects of immune activation not reflected when measuring gene abundances^25–27^. In several cases, infection induced switches in the most abundant equivalence class mapping to distinct transcripts, potentially altering coding regions or regulatory elements without modifying the overall gene output. Such isoform switches may reshape protein domains or regulatory motifs that influence signaling strength or mRNA stability, thereby modulating immune activation while maintaining core antiviral gene expression programs^3,4,13,56,76^. Our results underscore the importance of isoform-level analyses for a comprehensive understanding of immune responses and establish transcript usage as a complementary mechanism to gene expression.

Beyond infection-induced isoform remodeling, we further show that genetic ancestry shapes transcript usage variation during immune activation. Although ancestry is known to influence transcript usage variation^25,30–34,38^, its impact on isoform regulation during immune activation has remained largely unexplored beyond isolated cell types^25,36^. Population-associated differences in isoform usage are pervasive across lineages in baseline and infection contexts, yet predominantly lineage-specific and, in many cases, apparent only upon viral stimulation. Consistently, we identified population-specific differences in transcript usage response to infection, suggesting that populations differ not only in their isoform composition at baseline but also on how transcript usage programs are engaged during antiviral response. Such differences may influence immune function by modulating signaling pathways through receptor composition changes, as illustrated by the population-dependent isoform shifts observed for the CD8+ T cell exhaustion receptor regulator *KLRC1*^*67,68*^ upon IAV infection (Fig. 4b). Together, our findings indicate that population diversity shapes antiviral immunity at the level of transcript usage across immune lineages, adding another regulatory dimension beyond gene expression differences^42,43^.

Genetic contributions are expected to underlie part of the population differences in transcript usage^25,36,38,40^; however, we find that their influence differs between baseline and stimulated conditions, and that, under stimulation, these effects are virus-specific. In line with this, population DTU genes are enriched for TU-QTLs, indicating that ancestry-associated isoform differences frequently map to genetically regulated loci. Notably, these genetic effects vary across immune contexts. While many TU-QTLs are shared between baseline and stimulation contexts, a considerable fraction acts only upon viral stimulation (14–43% of TU-QTLs across immune lineages), which could explain the presence of genotype-by-environment interactions that remain silent at baseline. Moreover, a subset of these reTU-QTLs is specific to either IAV and COV, suggesting that genetic architecture of transcript usage host response differs between pathogens^25,28,41^. Collectively, these findings suggest that ancestry-associated transcript usage differences during viral stimulation arise from a combination of stable and stimulation-specific genetic effects and may, in part, be pathogen-dependent. This underscores the need to examine population differences across diverse immune challenges, as regulatory variation reflects the interaction between genetic background and environmental context. Extending these analyses will be necessary to determine whether pathogen-specific regulatory architectures represent a general feature of immune activation.

Some limitations of this study should be considered. First, our approach to model DTU does not account for overall gene expression levels, and may therefore be less precise for lowly expressed genes^77^. Second, our framework does not allow the identification of multiple independent genetic effects on transcript usage at the same locus, which may limit certain downstream analyses such as colocalization with GWAS signals^25,28,40^ or mediation analyses linking genetic variants to transcript usage variation^25,38,43^. Third, the largest scRNA-seq datasets under viral stimulation^42,43^ include only male donors, leaving the impact of sex in transcript usage variation during immune response unaddressed. Given well-established sex-specific differences in immune responses^78,79^, studies in sex-balanced cohorts will be necessary to determine whether transcript usage regulation exhibits sex-dependent patterns. Finally, our analyses are restricted to two human populations; extending this work to more diverse ancestries will be essential to broaden the landscape of immune transcript usage variation during antiviral responses. Despite these limitations, our study reveals how viral infection reshapes transcript usage programs across immune cell types and how genetic variation contributes to this process. These findings highlight isoform regulation as an important component of antiviral immune responses and motivate future studies exploring transcript usage across different pathogens, immune cell states, and human populations.

## Methods

### Ethical statement

This study was conducted in compliance with all relevant ethical guidelines. All analyses were based solely on publicly available datasets. No new human participants were recruited, no additional biological samples were collected, and no identifiable personal data were accessed. Ethical approval and written informed consent were obtained in the original studies.

### Data overview

We analyzed scRNA-seq data from PBMCs (769,534 cells) in resting conditions and after 6h stimulation with IAV and COV viruses from Aquino et al.^43^, including 160 individuals of African and European ancestry from the EVOIMMUNOPOP cohort^66^. In addition, we analyzed scRNA-seq (255,731 cells) and bulk RNA-seq data from PBMCs in resting conditions and after 6 hour stimulation with IAV from Randolph et al.^42^, comprising 89 male individuals of mixed African and European ancestry.

### Generation of a filtered gene annotation with bulk RNA-seq data

Filtering lowly expressed transcripts and genes from the reference annotation has been shown to increase the statistical power of DTU analyses in bulk RNA-seq by reducing the multiple-testing burden^80^. We therefore generated a filtered human transcriptome annotation (GENCODE v42) using bulk RNA-seq data from PBMCs generated by Randolph et al.^42^.

Raw sequencing reads from bulk RNA-seq data were processed as follows. First, FASTQ files were trimmed to remove adapters and low-quality bases using Trim Galore (v0.6.6) with default parameters. Transcript abundances were then estimated using kallisto quant^47^ (v0.48.0) using a transcriptome index constructed with *kallisto index* from GENCODE v42 human transcript sequences along with influenza A virus (H1N1) transcripts obtained from NCBI (accession GCF_001343785.1). The *--make-unique* option to handle duplicate transcript names. Transcripts were retained if they met the following expression criteria: transcripts per million > 1 (normalized) and estimated counts > 5 (unnormalized) in at least 20% of samples (N = 36). Transcripts originating from genes located in pseudoautosomal regions were excluded. After filtering, the final reference annotation comprised 33,787 transcripts corresponding to 15,779 genes.

### Equivalence class quantification of scRNA-seq data

Single-cell data were quantified using EC-based methods from the kallisto bustools workflow^47^. We first constructed a transcriptome index using *kallisto index*^*47*^ based on the filtered annotation described above. Next, for each sequencing library, raw FASTQ files were processed with *kallisto bus* from the bustools suite^81^(v0.41.0), setting the *--technology* flag to *10xv3* to match the input data format, to obtain the EC counts per cell (*EC × cell* count matrix). Barcode error correction was performed using *bustools correct* with a whitelist specific to the data type obtained from 10X Cell Ranger^82^ (v7.0.0) software. BUS files were subsequently sorted using *bustools sort*. Next, *gene x cell* and *EC x cell* count matrices were generated with *bustools count*, with and without the --genecounts flag, respectively. Transcript identifiers were assigned to equivalence classes based on their position in the transcriptome index. Finally, to estimate transcript-level abundances (*Transcript x cell* count matrices) at single-cell resolution, we applied the EM algorithm using *kallisto quant-tcc* on the *EC × cell* count matrices..

### Aquino et al. data processing

#### scRNA-seq preprocessing and pseudobulk generation

*Gene × cell* and *EC × cell* count matrices were generated using bustools^81^ for each of the scRNA-seq libraries using the same kallisto–bustools workflow described above, with the transcriptome index extended to include COV transcripts obtained from the NCBI Virus portal (accession PP405601.1). Matrices were imported into the R package Seurat (v5.0.2)^83^, and cells were assigned to individual donors using the cell- and donor-level metadata provided in Aquino et al.^43^. We filtered genes and ECs present in fewer than 10 cells, and filtered out cells with <500 genes, <800 and >25,000 counts, and >20% mitochondrial reads. Cells filtered at the gene level were also excluded from the EC count matrices to ensure consistency. We then merged each gene and EC count matrices by immune lineage. This resulted in five lineage-specific (CD4+ T cells, CD8+ T cells, monocytes, NK cells, and B cells) count matrices each containing baseline, IAV-, and COV-stimulated cells and the union of genes or ECs detected across libraries. For each immune lineage, gene and EC counts were aggregated by donor and condition using sparseSums function from textTinyR package (v.1.1.8). Counts from technical replicates were averaged using the arithmetic mean. Finally, genes and ECs with a mean counts-per-million value below 1 across all conditions (baseline, IAV, and COV) were excluded from further analyses.

### Randolph et al. data processing

We analyzed the Randolph et al.^42^ dataset for two distinct purposes: (i) to generate a bulk RNA-seq–filtered gene and transcript annotation, as described above; and (ii) to evaluate the reliability of transcript abundance estimation in 3′ scRNA-seq data by comparing transcript estimates derived from bulk RNA-seq and donor-level pseudobulk scRNA-seq samples.

#### Genotype preprocessing and cell demultiplexing

To demultiplex cells from each scRNA-seq library in Randolph et al.^42^ into individual samples, we first processed matched genotype data. Whole genome sequencing FASTQ files were aligned against the human reference genome (GRCh38.p13) using Burrows-Wheeler Aligner (BWA-MEM v0.7.17)^84^. Resulting BAM files were sorted, duplicate reads were marked, and read groups were added using samtools^85^. Variant calling was performed separately for each of the 22 autosomes using FreeBayes^85,86^. Per chromosome variant call format (VCF) files were filtered to retain variants with minor allele frequency (MAF) > 0.05, QUAL > 30, missingness < 10%, and read depth between 2 and 50. Filtered variants were subsequently imputed using Beagle^87^. Finally, chromosome-level VCFs were concatenated using bcftools^85^ resulting in a processed genotype data. For demultiplexing, raw scRNA-seq FASTQ for each library were processed with Cell Ranger^82^ (v7.0.0, 10X Genomics) using a reference transcriptome comprising GENCODE v42 human transcripts and influenza A virus (H1N1) transcripts (GCF_001343785.1 NCBI accession). Cells were then demultiplexed into donors using Souporcell^82,88^ with the processed genotype data, and doublets were excluded based on Souporcell assignments.

#### scRNA-seq preprocessing and pseudobulk generation

*Gene × cell, and transcript × cell* count matrices were generated as described above and imported into Seurat (v5.0.2)^83^. Cells were filtered to retain those with 200–5,000 detected genes and <10% mitochondrial reads, and genes or transcripts expressed in fewer than 10 cells were excluded. Cells filtered during gene-level filtering were also excluded from the EC and transcript count matrices to ensure consistency across analysis. *Transcripts x cell* matrices were merged across libraries using RowMergeSparseMatrices function from Seurat (v5.0.2)^83^. Finally, transcript counts were aggregated at the donor level by pseudo-bulking using sparseSums function from textTinyR (v.1.1.8).

### Comparing transcript abundances obtained from bulk RNA-seq and 3’ scRNA-seq data

To evaluate whether transcript abundances can be reliably inferred from 3’ scRNA-seq data, we compared transcript abundance estimates derived from bulk RNA-seq and donor-matched single-cell pseudobulk profiles using data from Randolph et al.^42^. Specifically, we computed Pearson correlations between transcript abundances estimated from bulk RNA-seq PBMC samples and those inferred from donor-level pseudobulk profiles of 3′ scRNA-seq data for the 12,136 transcripts detected in both datasets. We considered significant correlations under FDR<0.05.

### Transcriptome complexity across single-cells

To quantify transcriptome complexity at the single-cell level in the Aquino et al.^43^ dataset, we defined a metric based on the number of ECs expressed per gene, termed ‘cell EC diversity’. Specifically, for each immune lineage *EC x cell* count matrix, we first derived a *gene × cell* matrix in which each entry represents the number of ECs expressed (count > 0) for a given gene in a given cell. ‘Cell EC diversity’ was then calculated for each cell as the mean number of expressed ECs per gene across all genes, restricting the analysis to genes with at least two annotated transcripts in the filtered transcriptome reference. Significant differences in cell EC diversity between baseline and stimulated conditions (IAV, COV) were assessed using the Wilcoxon rank-sum test (FDR<0.05). To control for variation in sequencing depth between cells, we selected, for each immune lineage, the top 75% of cells by UMI count and downsampled them to a common minimum UMI depth using downsampleMatrix from scuttle^89^. ‘Cell EC diversity’ was then recalculated on the downsampled data and compared between baseline and stimulated cells.

### Differential gene expression analysis

To conduct differential gene expression analysis we relied on dream^90^, which fits linear mixed models (LMM) and allows simultaneous modeling of fixed and random effects. All analyses were performed independently for each immune lineage using the donor-aggregated expression data described above. For each gene, a LMM was fitted using the dream function.

#### Infection-associated differential gene expression

To identify infection-associated DEGs, we performed pairwise comparisons between baseline and each viral stimulation. Specifically, gene expression pseudo-bulked counts of infected and baseline samples were grouped in a pairwise manner (e.g. IAV-Baseline, COV-Baseline). Then, for each gene in each lineage we fit the following model:

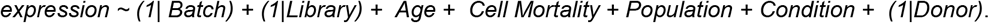

Because technical replicates in the Aquino et al.^43^ dataset belonged to different sequencing libraries, library identifiers from paired replicates were combined into a composite covariate to properly account for library effects. Genes with |logFC|>0.5 and FDR<0.01 (Benjamini-Hochberg correction) were considered infection DEGs.

#### Population-associated differential gene expression

Using the same donor-aggregated expression data and modeling strategy, we next tested for population-associated gene expression differences in each immune lineage and baseline or stimulation condition separately (e.g. CD4+ T-Baseline). For each gene, we fitted the following model:

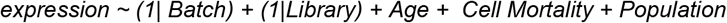

To further adjust for interindividual variation in immune cell composition within each lineage we included the relative abundances of cell type subsets. For example, within NK cells, the proportion of CD56^brt^, CD56^dim^ and memory NK cells were included:

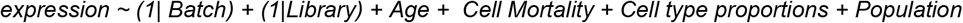

For CD4+ T cells, we excluded the smallest subset (CD4+ T regulatory cells) due to a rank-deficient design matrix due to perfect multicollinearity and model failure. We considered genes with |logFC|>0.2 and FDR<0.01 to be population DE.

### Equivalence class ratio calculation

To model transcript usage differences upon viral infection and between populations, we quantified EC relative abundances from EC count matrices. EC counts were first grouped according to the biological comparison of interest. Specifically, for infection DTU analyses, EC counts were organized into paired contrasts between baseline and stimulated conditions (e.g. IAV–Baseline or COV–Baseline). For population-associated analyses, EC counts were stratified by condition (Baseline, IAV, or COV) to allow population effects to be tested within each context. Within each group, EC relative abundances were computed as the proportion of each EC count relative to the total EC counts for the corresponding gene. Relative abundance estimation and filtering were performed using the prepare.trans.exp function from *sQTLseekeR2*^*37*^. ECs were required to have more than one count across all samples (min.transcript.exp = 1), samples required gene counts greater than five (min.gene.exp = 5), and genes were required to be present in at least 80% of samples (min.prop = 0.8). A minimum EC dispersion of 0.01 was applied (min.dispersion = 0.01). Genes retaining only a single EC after filtering were excluded from downstream analyses.

### Differential transcript usage analysis upon viral stimulation

To identify viral infection-associated transcript usage differences, we developed a novel computational framework for transcript usage using paired data based on the multivariate non-parametric test implemented in the MANTA R package^51^. Specifically, for each donor, we subtracted the EC ratios of infected and baseline samples to generate EC ratio change vectors (e.g. Donor_1 IAV_-Donor_1Baseline_). These EC difference vectors were then jointly modelled at the gene level. For each gene within each immune lineage and viral response (e.g. CD4+ T IAV-Baseline), we fitted the model:

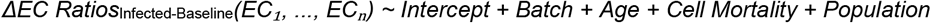

Within this framework, the Intercept term captures the systematic shifts in EC relative abundances between infected and baseline conditions. Genes with significant Intercept after Benjamini–Hochberg correction, (FDR < 0.05) were classified as infection DTU genes. Sequencing library was not included as a covariate in DTU models, as the large number of library levels led to model convergence failures.

Because EC relative abundances are bounded between 0 and 1, modeling differences between two EC ratio matrices can induce strong correlations among EC vectors, potentially leading to rank-deficient design matrices. When such rank deficiency was encountered, we randomly removed one EC from the model. For genes with only two ECs exhibiting complete anticorrelation, a single EC vector was modeled.

### Population-specific differential transcript usage in response to infection

To identify population-specific transcript usage responses to infection, we applied the same modeling framework explained in the previous section while additionally accounting for population differences in cell composition. Specifically, we included as covariates the lineage-specific cell type subset abundances in the infected condition. For each gene within each immune lineage and viral response (e.g. CD4+ T IAV-Baseline), we fitted the model:

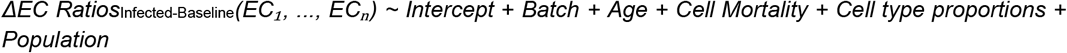

In this extended model, the Population term captures population-by-infection interaction effects. Genes with a significant Population term (FDR < 0.05) were classified as population differential response transcript usage genes.

### Differential transcript usage analysis between populations

To identify genes with population-associated transcript usage differences, we used the non-parametric multivariate test implemented in the MANTA R package^51^ separately within each immune lineage and condition (e.g. CD4+ T-Baseline). For each gene, we fitted the following model:

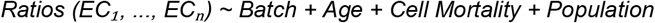

We also applied another model including lineage-specific cell type subset abundances as covariates, as described in the differential gene expression analyses:

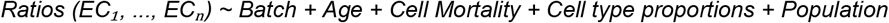

Genes with a significant *Population* term after Benjamini–Hochberg correction (FDR < 0.05) were classified as population DTU genes.

### DTU effect size

As a measure of DTU effect size, we used the maximum absolute difference (MD value) in mean EC relative abundance between groups, either infected versus baseline samples or between populations. MD values were computed using the md.trans function from *sQTLseekeR2*^*37*^ R package.

### Isoform switch detection in DTU genes

To identify genes exhibiting a switch in their dominant EC, we first calculated the mean EC relative abundance within each group of samples (e.g. by condition for infection DTU or by population for population DTU). A change in the most abundant EC between groups was defined as an EC switch. EC switches were further classified into two categories: (i) isoform switches, where the switching ECs correspond to non-overlapping sets of transcripts (e.g. EC_1_ {t_1_} versus EC_2_ {t_2_}); and (ii) non-isoform switches, where the switching ECs partially overlap in transcript composition (e.g. EC_1_ {t_1_, t_2_} versus EC_2_ {t_2_}).

### Pathway enrichment analysis

To identify biological pathways enriched among gene sets exhibiting transcript usage differences, we performed overrepresentation analyses with clusterProfiler^91^ (v3.20) using the org.Hs.eg.db (v3.20) database, considering all tested genes as the background universe. Gene Ontology (GO) terms containing at least five genes were retained (minGSSize = 5), and significance was assessed using a Benjamini–Hochberg–adjusted false discovery rate (FDR < 0.05; pvalueCutoff = 0.05, pAdjustMethod = “BH”). To summarize and reduce redundancy among enriched GO terms, we used rrvgo^58^ (v.1.10.0), using the calculateSimMatrix function (method=“Rel”) and clustering terms using the reduceSimMatrix function with similarity threshold = 0.7.

### Assigning genomic coordinates to equivalence classes

To improve the interpretability of our transcript usage analysis, we developed a strategy to assign ECs to genomic coordinates. Each EC represents the set of transcripts sharing sequence regions from which a sequencing read could have originated from^47–49,92^ (Fig 1.b). Based on this definition, we implemented a two-step procedure to map ECs to genomic regions (<u>Supplementary Fig. 23</u>). First, for each EC, we computed the intersection of the genomic coordinates of all transcripts it contains. For example, EC1 {t1} is assigned the coordinates of transcript t1; EC2 {t1, t2} is assigned the intersection of t1 and t2; and EC3 {t2} is assigned the coordinates of t2. Second, to obtain EC-specific, non-overlapping regions, we subtracted shared coordinates between ECs. Specifically, the intersected coordinates of EC2 {t1, t2} were subtracted from those of EC1 {t1} and EC3 {t2}, yielding mutually exclusive genomic segments. This ensures that no two ECs map to the same genomic region, consistent with the conceptual definition of ECs as distinct transcript segments. Importantly, coordinate assignment was performed considering all filtered ECs, not only those retained for downstream transcript usage testing.

We note that this coordinate-based approach provides only an approximation and does not capture the exact genomic sequences corresponding to each EC. ECs are defined by shared *k*-mer matches between query reads and reference transcripts^47,92^. This limitation is particularly relevant for 3′ scRNA-seq data, where coverage is biased toward transcript ends. As a result, ECs may be assigned to transcript regions that are not directly supported by read coverage, such as 5’ transcript ends. To facilitate accurate interpretation, we therefore recommend visualizing EC coordinates alongside read coverage profiles when examining specific loci.

To visualize read coverage alongside EC coordinates, we aligned Aquino et al., ^43^ scRNA-seq data to the GRCh38.p13 genome sequence using a reference index built with cellranger mkref (10X Cell Ranger^82^, v7.0.0) containing GENCODE v42 human transcripts, influenza A and SARS-CoV-2 virus transcripts.BAM files for each sequencing library were generated using cellranger count. Each BAM file was split by donor, immune lineage, and baseline or stimulated condition (e.g CD4+ T-Baseline-Donor_1_). To compute gene-level coverage across multiple donor-specific BAM files, we used samtools depth^85^ while filtering low quality reads using the -q 30 parameter. Integrated visualizations of EC coordinates and read coverage were generated using the Gviz R package^93^. Specifically, transcripts, ECs, and coverage tracks Transcripts, ECs and coverage were visualized GeneRegionTrack, HighlightTrack and DataTrack functions respectively.

### Permutations of transcript usage analyses

To evaluate the robustness of our framework and estimate its false positive rate, we performed permutation analyses by randomly shuffling sample labels and re-running the corresponding transcript usage models. For infection DTU, where EC ratio differences are modeled between paired conditions (e.g., IAV–Baseline), we randomly permuted condition labels across donors within each immune lineage, generating mismatched infected–baseline pairs (e.g. Donor_1 Infected_-Donor_1 Baseline,_ Donor_2 Baseline_-Donor_2 Infected_) for each immune lineage. This procedure was repeated 100 times, and genes with a significant Intercept term after Benjamini–Hochberg correction (FDR < 0.05) were considered false positives. For population × infection interaction analyses, we applied the same permutation strategy while fitting the interaction model, defining genes with a significant Population term (FDR < 0.05) as false positives in 100 permutations. Finally, to assess false positives in population DTU analyses, donor labels were shuffled within each lineage and condition (e.g. CD4+ T cells – Baseline) and the analysis repeated 100 times; genes with FDR < 0.05 for the Population term were counted as false positives.

### Mapping the genetic determinants of transcript usage

To identify genetic variants associated with transcript usage variation (TU-QTLs), we modelled EC ratios across genotype groups using the multivariate non-parametric framework implemented in the *sQTLseekeR2*^*37*^ Nextflow^94^ pipeline (sqtlseeker2-nf, v1.1.1). We started from a set of 13.6 million SNPs that had been filtered, phased, and imputed, provided by Aquino et al.^43^. Genotypes were converted to 0/1/2 dosage format using vcftools^95^ (v0.1.16) with the --*012* option. For each immune lineage and condition (e.g. CD4+ T-Baseline), EC count matrices were provided as input to the sqtlseeker2-nf pipeline, applying filtering criteria analogous to those used in the DTU analyses. Genes located within the HLA super-locus^96^ were excluded, consistent with previous studies^25,40^. To correct for population structure, five genotype principal components (PCs) calculated using PLINK v2.0^97^ were added into the model. In addition, five PCs derived from the EC ratio matrix were incorporated to account for global transcript usage variation. To account for cell composition differences between individuals, we included the subcell type abundances of each immune lineage as covariates. For each gene–SNP pair within a given lineage and condition, we fitted the following model:

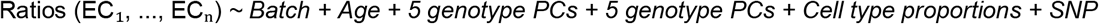

Significant TU-QTLs were identified using a Benjamini-Hochberg FDR threshold of 0.01 (--type_fdr=BH and --fdr=0.01). To detect stimulation-specific TU-QTLs (response TU-QTLs), we considered gene-SNP pairs that were significant upon viral stimulation (FDR_Infected_<0.01) but not significant at baseline (p_Baseline_ > 0.01).

### Overlap with external datasets

To validate our findings, we overlapped our results with those obtained in different independent studies that mapped splicing differences in the immune system using Fisher’s exact test (FDR<0.05). First, we used the study by Rotival et al.^25^, which quantified splicing via percentage-spliced in (PSI) in a bulk RNA sequencing of human primary monocytes at basal state and after stimulation with different bacterial and viral ligands in 200 individuals of African (N=100) and European (N=100) descent. This study reported alternative splicing differences upon infection, between human populations and mapped splicing QTLs. We overlapped our monocyte DTU genes in response to IAV and COV infection with those obtained by Rotival et al.^25^ after IAV stimulation. Similarly, we compared our monocyte population DTU genes with their population splicing differences both in resting conditions, and our population DTU genes in infected samples (IAV, COV) with their IAV-stimulated data. We also evaluated overlap between our TU-QTL genes and the splicing QTL genes identified in Rotival et al.^25^, separately for baseline and infection conditions. Additionally, we compared our TU-QTL results obtained in baseline with those from Garrido-Martín et al.^37^ where sQTLs were mapped across healthy tissues in the GTEx v8 dataset using sQTLseekeR2 method. Specifically, we overlapped our TU-QTLs and TU-QTLs genes with the sQTLs and sGenes identified in EBV-transformed lymphocytes and whole blood tissues. Finally, we assessed overlap between our baseline TU-QTLs and TU-QTL genes and the sQTLs and sGenes recently mapped across immune cell types in Asian individuals using 5′ scRNA-seq data^40^.

## Data availability

All data analysed in this study are publicly available. The scRNA-seq and genotype data from Aquino et al.^43^ are available through the Institut Pasteur data repository OWEY at https://doi.org/10.48802/owey.e4qn-9190 and https://doi.org/10.48802/owey.pyk2-5w22, respectively. Single-cell and bulk RNA-seq data from Randolph et al.^42^ were downloaded from GEO^98^ database (GSE162632). Whole-genome sequencing data for the same study were retrieved from SRA (PRJNA736483). Summary statistics used to compare our results with external datasets were obtained from the original publications ^25,37,40^.

Supplementary tables derived from the analysis conducted in this paper are deposited in the following link: https://drive.google.com/drive/folders/1WCI7FhCU8dL5vHM7Wwzve1zERghhRhSd?usp=sharing

## Code availability

All the custom scripts developed for this study have been made openly accessible and can be found on the GitHub repository at: https://github.com/rubenchazarra/2026_infection_transcript_usage. By visiting this repository, researchers and interested individuals can access and use the scripts for their own analysis or to replicate the study’s findings.

## Acknowledgements

R.C.G was supported by a predoctoral *Formación de Personal Investigador (FPI)* fellowship from the *Ministerio de Ciencia, Innovación y Universidades (MCIN)* and the *Agencia Estatal de Investigación (AEI)* (PRE2020-092510) funded by MCIN/AEI. A.R.C. was supported by a predoctoral *FPI* fellowship from the *MCIN* and the *AEI* (PRE2019-090193). M.S.R. was supported by a predoctoral *AGAUR-FI Joan Oró* fellowship from the *Secretaria d’Universitats i Recerca, Departament de Recerca i Universitats, Generalitat de Catalunya*, and the *European Social Fund Plus* (FI-3 2024-0065). This project was supported by grant PID2019-107937GA-I00 and M.M. was supported by grant RYC-2017-22249, both funded by MCIN/AEI/10.13039/501100011033. This project was supported, by the *AGAUR-SGR* grant from the *Secretaria d’Universitats i Recerca, Departament de Recerca i Universitats, Generalitat de Catalunya* (2021 SGR 01577), which co-financed this work, and by the *European Social Fund* (“Investing in Your Future”) and the Chan Zuckerberg Initiative on the Ancestry Networks for the Human Cell Atlas REF: AN-0000000078.

We thank Winona Oliveros, José Miguel Ramírez, Pau Clavell-Revelles, and Fairlie Reese for their valuable feedback during the development of this work. We thank Martin Hemberg and Jae-Won Cho for helpful discussions at the early stages of the project on strategies to quantify transcript usage in end-biased scRNA-seq data. We also thank Mark Robinson for valuable input on differential transcript usage modelling in single-cell data and feedback on the publication. Finally, we thank Yann Aquino and Maxime Rotival for helpful discussions about the dataset and differential splicing analyses accounting for cell composition differences between individuals. We dedicate this work to the memory of Miquel Calvo, whose insight and dedication greatly contributed to this study.

## Declaration of generative AI and AI-assisted technologies in the writing process

During the preparation of this work, the authors used ChatGPT to enhance the clarity and language of the text. After using these tools, the authors reviewed and edited the content as needed and take full responsibility for the content of the publication.

## Author contribution

R.C.G. designed, performed, and supervised all analysis.

A.R.C. and M.S.R. advised in data analysis and supervised all analysis.

I.M.P. led data analysis related to transcript usage QTLs (Figure 3).

M.C. and F.R. assisted in statistical analysis design and implementation

D. G-M. strongly contributed to the design and implementation of statistical analyses and provided methodological guidance.

M.M. conceived the study and supervised all analysis. R.C.G., A.R.C, M.S.R. and M.M wrote the manuscript.

