## Supplementary Figures for "Single-cell atlas of transcript usage remodelling in antiviral immune responses across human populations"

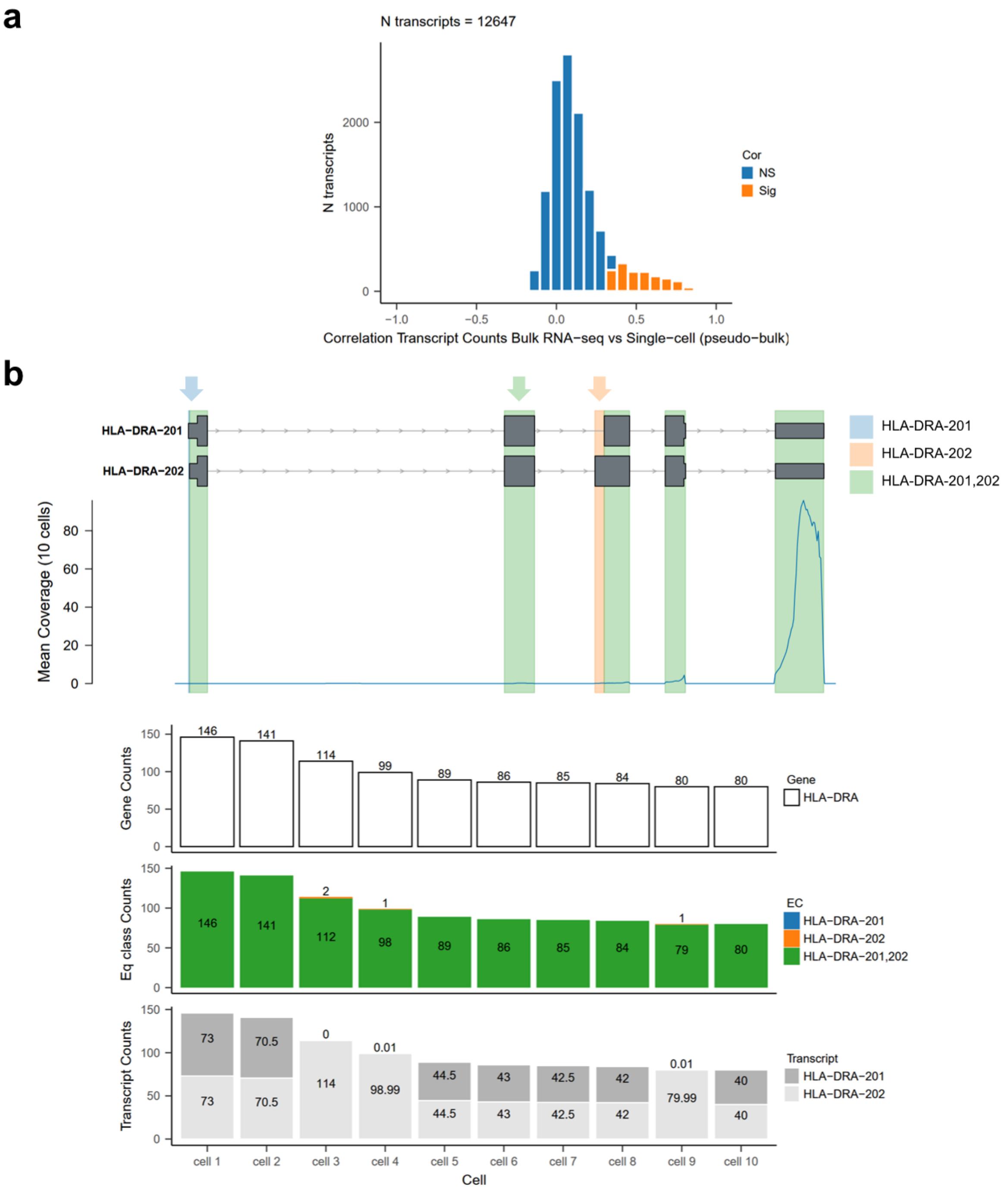


**Supplementary Fig. 1. Probabilistic estimation of transcript abundances in 3' single-cell RNA sequencing (scRNA-seq) data is unreliable due to the inherent transcript end coverage bias. a)** Distribution of Pearson correlation coefficients per transcript (N=12,136) between transcript abundance estimates obtained from bulk RNA-seq PBMCs quantified with kallisto[1] and from a pseudo-bulk of all PBMCs from a donor-matched 3' scRNA-seq dataset quantified using bustools[2]. Matched data is from ref[3]. **b)** Example of ambiguous transcript abundance estimation in the *HLA-DRA* gene. Top: representation of the 2 *HLA-DRA* transcripts with UTRs shown as thin lines and coding sequences as thick lines. Colored vertical blocks highlight the transcript segments represented by each equivalence class (EC): the green EC (*HLA-DRA-201,202*) corresponds to the shared structure between *HLA-DRA-201* and *HLA-DRA-202,* the blue EC (*HLA-DRA-201*) corresponds to *HLA-DRA-201*-specific region in exon 1, and the orange EC (*HLA-DRA-202*) corresponds to *HLA-DRA-202*-specific region in exon 3. The coverage track represents the summed read coverage of the 10 cells with the highest *HLA-DRA* counts in one library in ref[3]. Bottom: barplot showing gene-level counts, EC-level counts, and inferred transcript abundances from the same 10 cells. In cells where reads map exclusively to the shared EC (HLA-DRA-201,202, green EC), transcript abundance estimation assigns counts equally between the two transcripts.

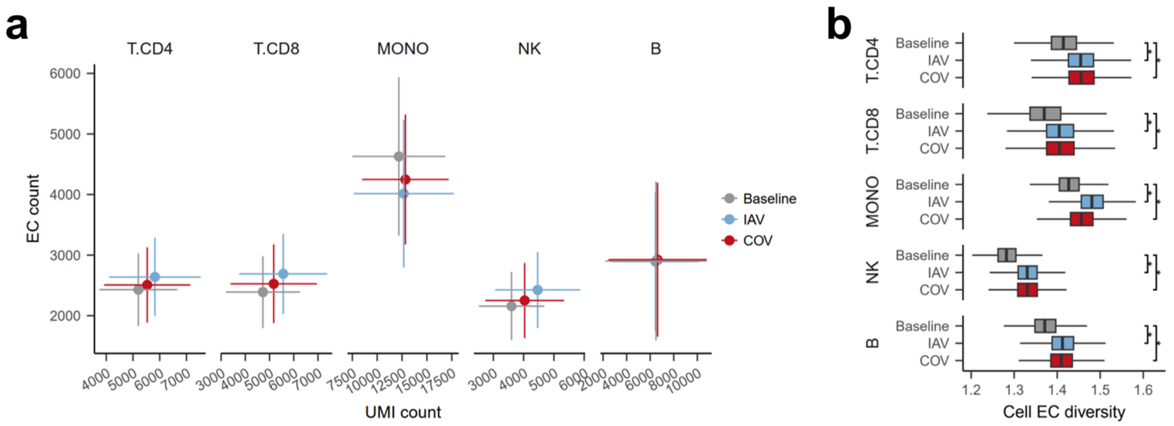


**Supplementary Fig. 2. Differences in sequencing depth do not explain the increase in transcriptome complexity upon viral stimulation. a)** Cells stimulated with influenza A virus (IAV) or SARS-CoV-2 (COV) exhibit higher total unique molecular identifier (UMI) counts and equivalence class (EC) counts per cell compared to baseline. For each immune lineage and condition, the mean number of UMIs (x-axis) and ECs (y-axis) across cells is shown; error bars indicate ±1 standard deviation. **b)** To control for differences in sequencing depth, we selected the top 75% of cells by UMI count within each immune lineage and downsampled to a common minimum UMI count. "Cell EC diversity" was then calculated as the mean number of ECs expressed per gene per cell, restricted to genes with ≥2 annotated transcripts. Each data point represents a single cell; outliers are not shown. After UMI normalization, virally stimulated cells (IAV, COV) consistently exhibit significantly higher cell EC diversity than baseline cells (Wilcoxon test; FDR< 2.2 x $10^{-16}$).

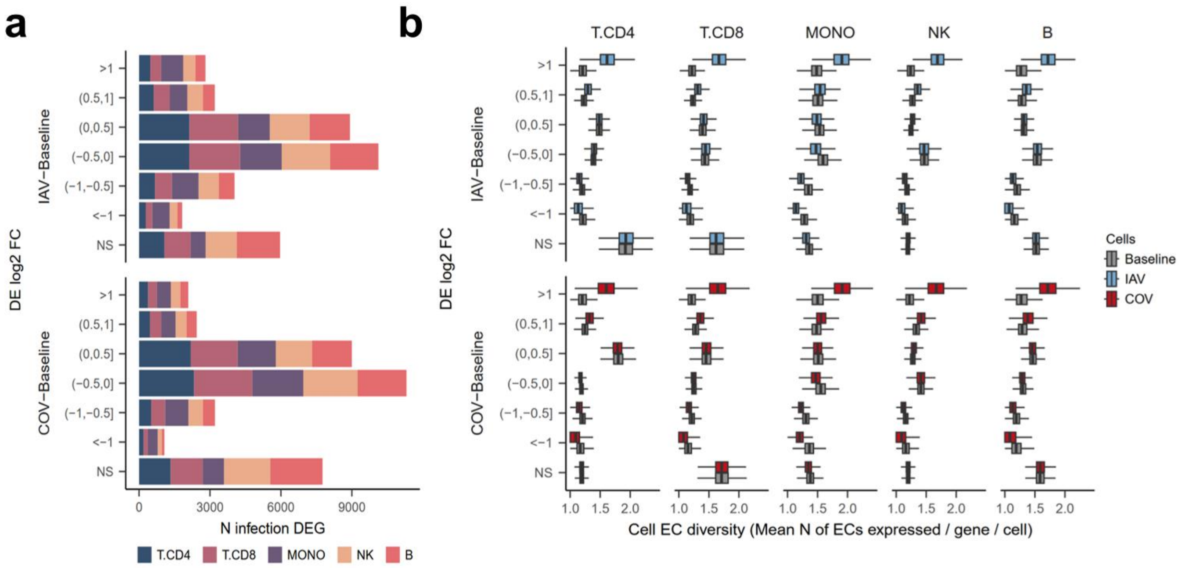


**Supplementary Fig. 3. Viral-associated increase in transcriptome complexity is largely driven by highly upregulated infection-responsive genes. a)** Number of infection-associated differentially expressed genes (DEG; FDR<0.01) stratified by their log2 fold change for each immune lineage and viral response (influenza A virus (IAV)-Baseline, SARS-CoV-2 (COV)-Baseline). NS, not significant (FDR>=0.01). **b)** "Cell equivalence class (EC) diversity", quantified as the mean number of ECs expressed per gene per cell, for the infection-associated DEGs shown in (a), comparing baseline , IAV- and COV-stimulated cells.

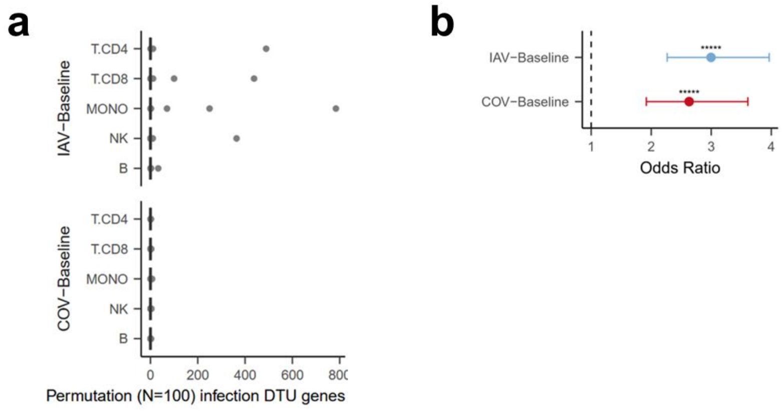


**Supplementary Fig. 4. Validation of infection-associated differential transcript usage (DTU) detection by permutation analysis and overlap with previous studies. a)** Permutation analysis for infection-associated DTU detection. Under the original (non-permuted) infection DTU analysis, in each viral response (influenza A virus (IAV)-Baseline and SARS-CoV-2 (COV)-Baseline), equivalence class (EC) differences were computed between matched infected and baseline samples from the same donor. (e.g. Donor$_{1\ \text{Infected}}$-Donor$_{1\ \text{Baseline}}$, Donor$_{2\ \text{Infected}}$-Donor$_{2\ \text{Baseline}}$). Here, instead, we randomly shuffled the condition labels across donors, such that EC ratio differences were calculated between mismatched infected and baseline samples (e.g., Donor$_{1\ \text{Infected}}$-Donor$_{1\ \text{Baseline}}$, Donor$_{2\ \text{Baseline}}$-Donor$_{2\ \text{Infected}}$). This procedure was repeated 100 times, and the infection DTU analysis was performed for each permutation. Boxplots show the number of significant DTU genes (FDR<0.05) detected per permutation (center line: median; box limits: upper and lower quartiles; whiskers: 1.5× interquartile range; points: outliers). **b)** Overlap with previously reported infection-associated splicing differences. Overlap between monocyte DTU genes identified in our study following IAV and COV infection and those reported in a bulk RNA-seq study of IAV-infected monocytes[4]. Enrichment is shown as odds ratio, with the dotted line indicating no enrichment (odds ratio = 1). Asterisks denote significance levels (* p < 0.01, ** p < 0.001, etc.).

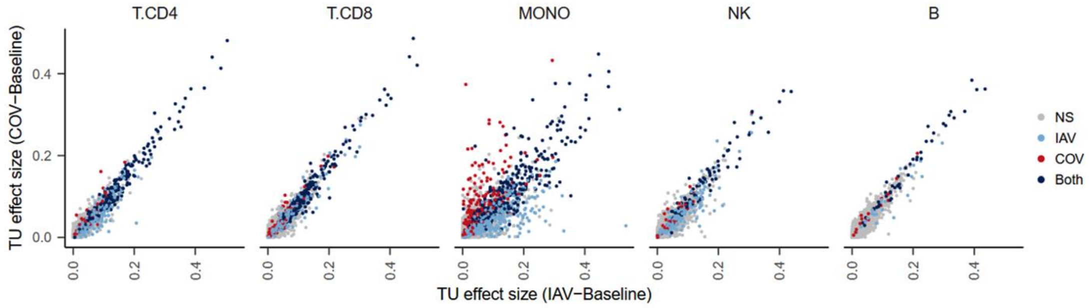


**Supplementary Fig. 5. Comparison of transcript usage responses to influenza A virus (IAV) and SARS-CoV-2 (COV) across immune lineages.** Each point represents a gene, plotted according to its differential transcript usage (DTU) effect size (MD value) in response to IAV (x-axis) and COV (y-axis) stimulation within each immune lineage. Colors indicate whether genes exhibit infection-associated DTU in response to IAV only, COV only, both viruses, or neither condition (NS).

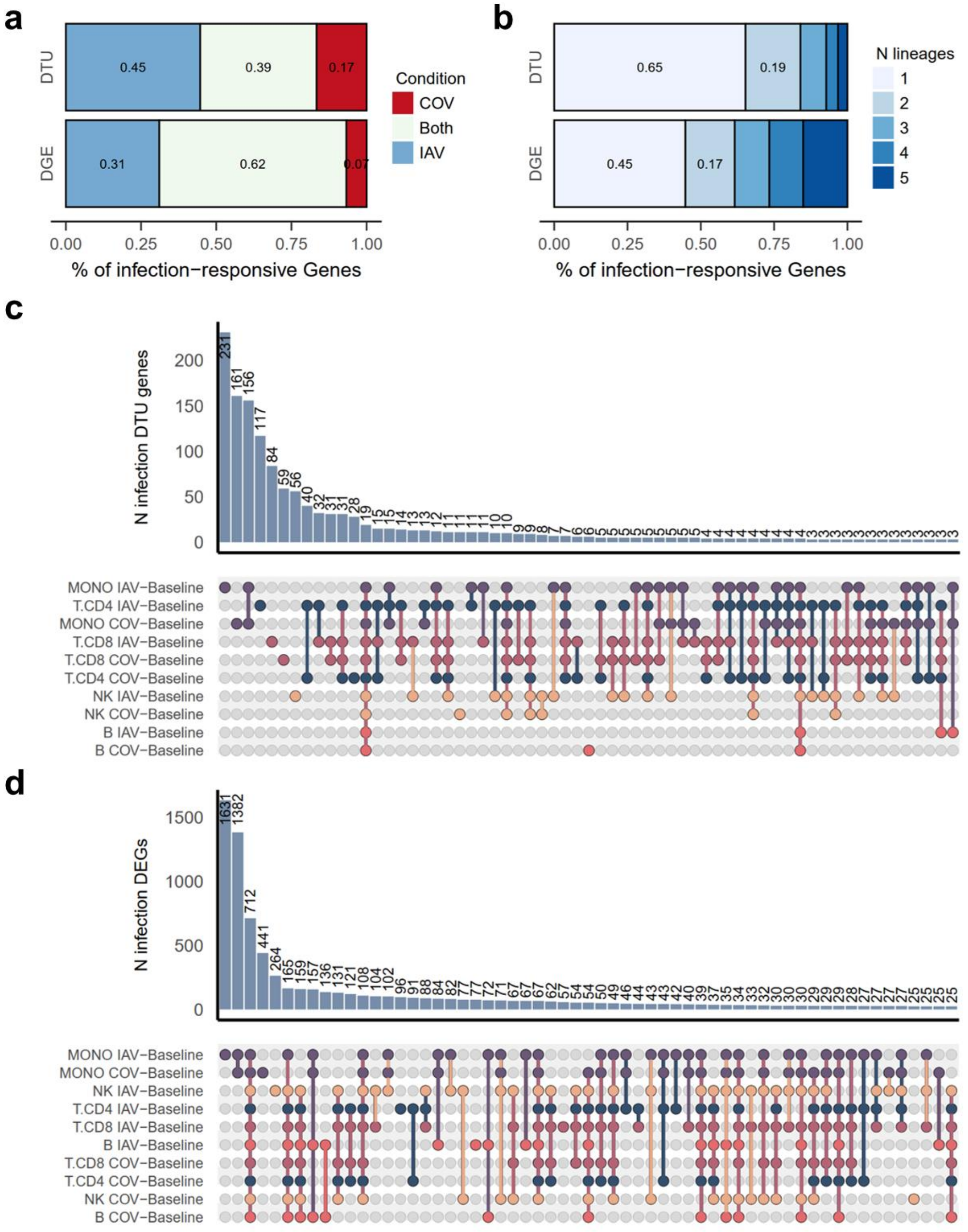


[Supplementary Fig. 6]. **Sharing patterns of infection-associated differential transcript usage (DTU) and differential gene expression (DGE) analyses.** **a)** Proportion of infection-responsive genes identified by DTU and DGE analyses that are specific to influenza A virus (IAV), specific to SARS-CoV-2 (COV), or shared between both responses. **b)** Proportion of infection-responsive genes identified by DTU and DGE analyses that are shared across increasing numbers of immune lineages. **c)** Upset plots showing the overlap of infection-associated DTU for gene sets containing more than 3 DTU genes. **d)** Upset plots showing the overlap of infection-associated DGE for gene sets containing more than 25 genes. In both plots, the analysis is restricted to genes tested in both responses within each immune lineage.

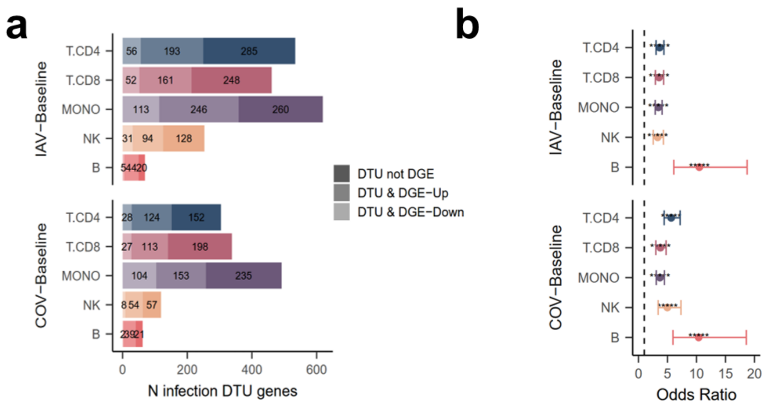


**Supplementary Fig. 7. Infection-associated differential transcript usage (DTU) is largely independent of overall gene expression changes. a)** Barplot of infection-associated DTU genes stratified by whether they are not differentially expressed upon infection (DTU not DGE), and differentially expressed and upregulated (DTU & DGE-Up) or downregulated (DTU & DGE-Down). **b)** Enrichment analysis between genes exhibiting DTU and DGE. The dotted line indicates no enrichment (odds ratio = 1). Asterisks denote significance levels (* $p < 0.01$, ** $p < 0.001$, etc.).

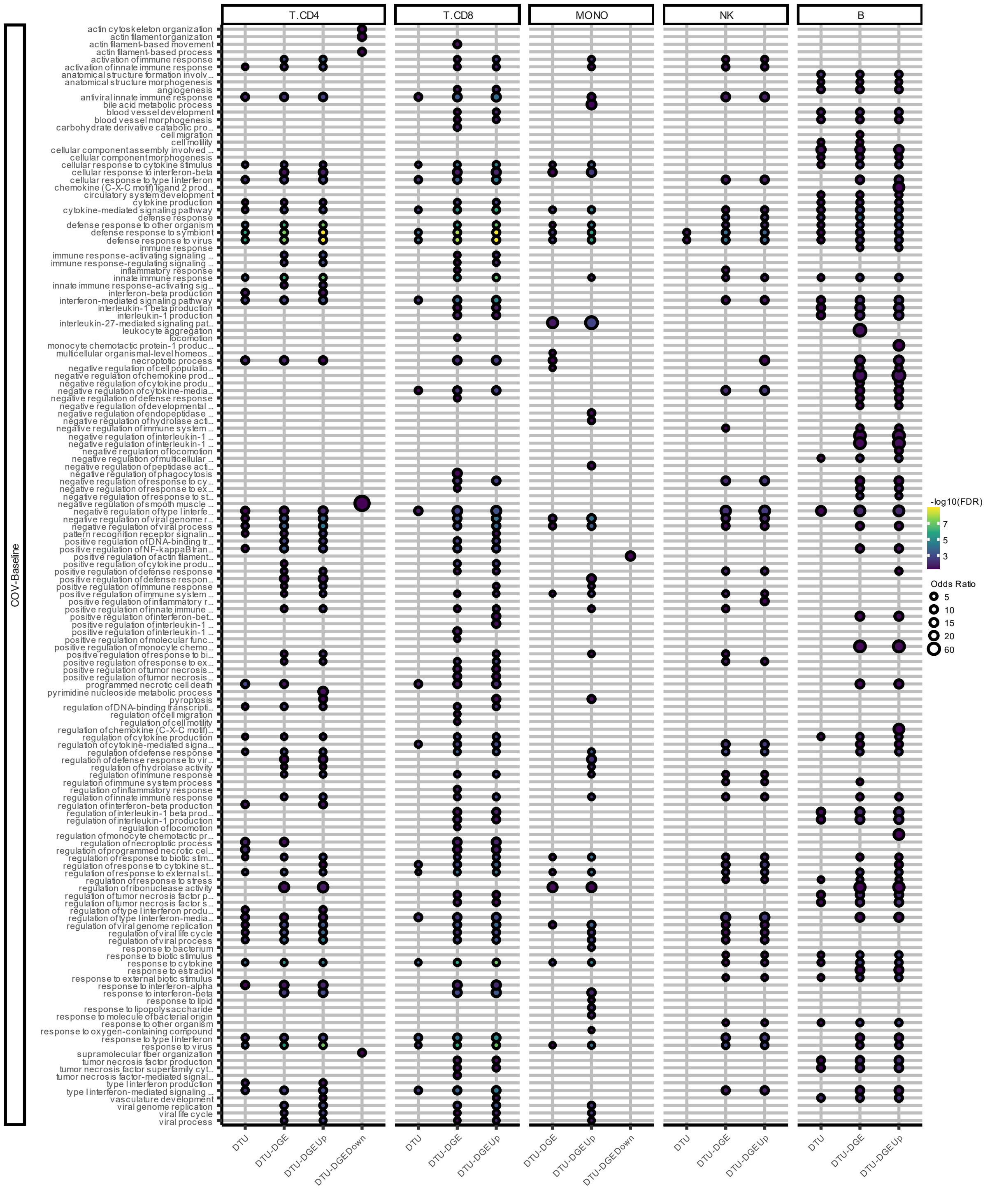


**Supplementary Fig. 8. Pathway overrepresentation analysis (ORA) of infection-associated differential transcript usage (DTU) genes in response to SARS-CoV-2 (COV) across immune lineages.** ORA was performed for different DTU gene sets: all DTU genes (DTU); DTU genes overlapping with differentially expressed genes (DTU–DGE); and DTU–DGE genes further stratified into upregulated (DTU–DGE Up) and downregulated (DTU–DGE Down) categories. Shown are Gene Ontology (GO) biological process terms with FDR < 0.05. Point color represents $-\log_{10}$(FDR), and point size reflects the enrichment odds ratio.

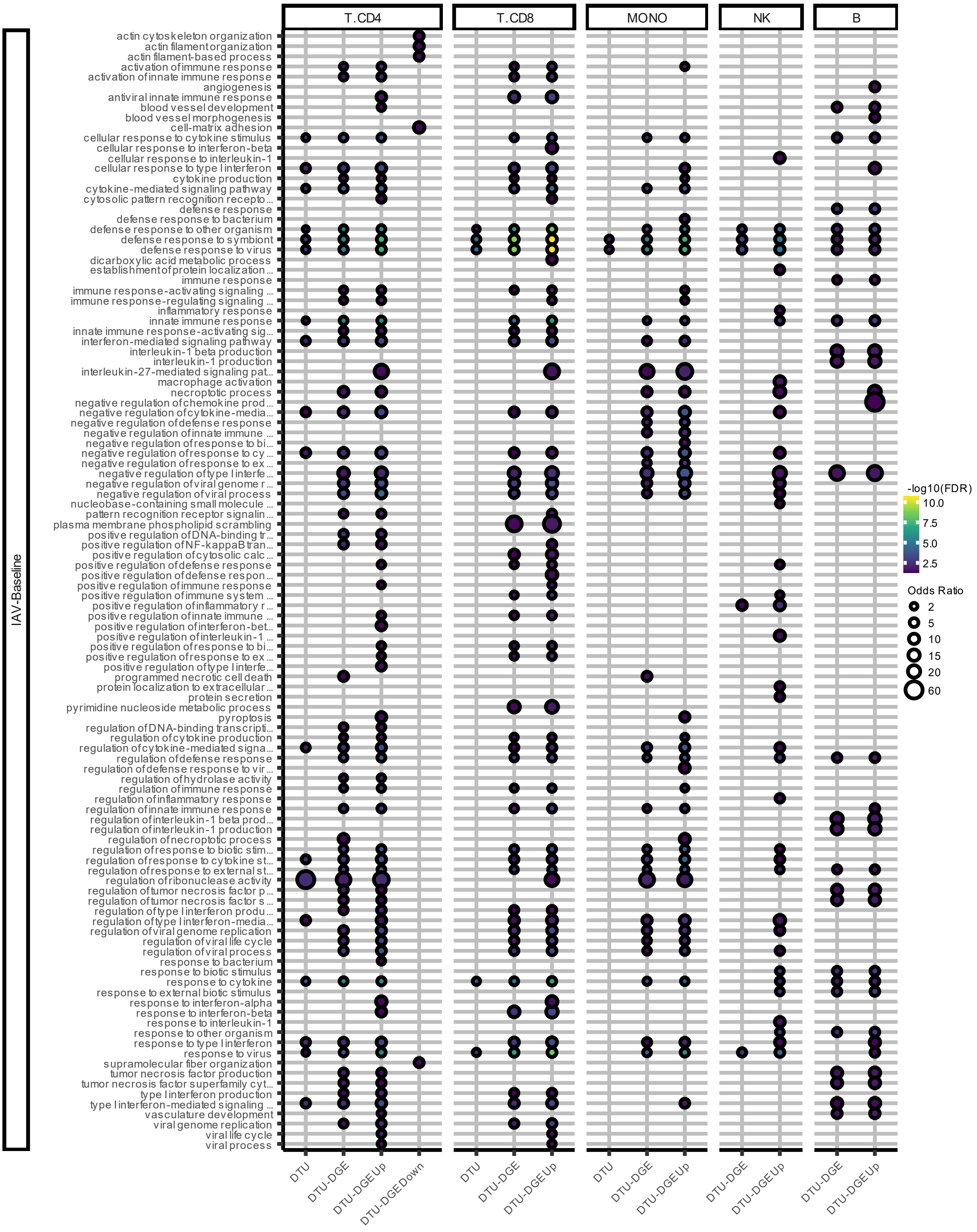


**Supplementary Fig. 9. Pathway overrepresentation analysis (ORA) of infection-associated differential transcript usage (DTU) genes in response to influenza A (IAV) across immune lineages.** ORA was performed for different DTU gene sets: all DTU genes (DTU); DTU genes overlapping with differentially expressed genes (DTU–DGE); and DTU–DGE genes further stratified into upregulated (DTU–DGE Up) and downregulated (DTU–DGE Down) categories. Shown are Gene Ontology (GO) biological process terms with FDR < 0.05. Point color represents $-\log_{10}$(FDR), and point size reflects the enrichment odds ratio.

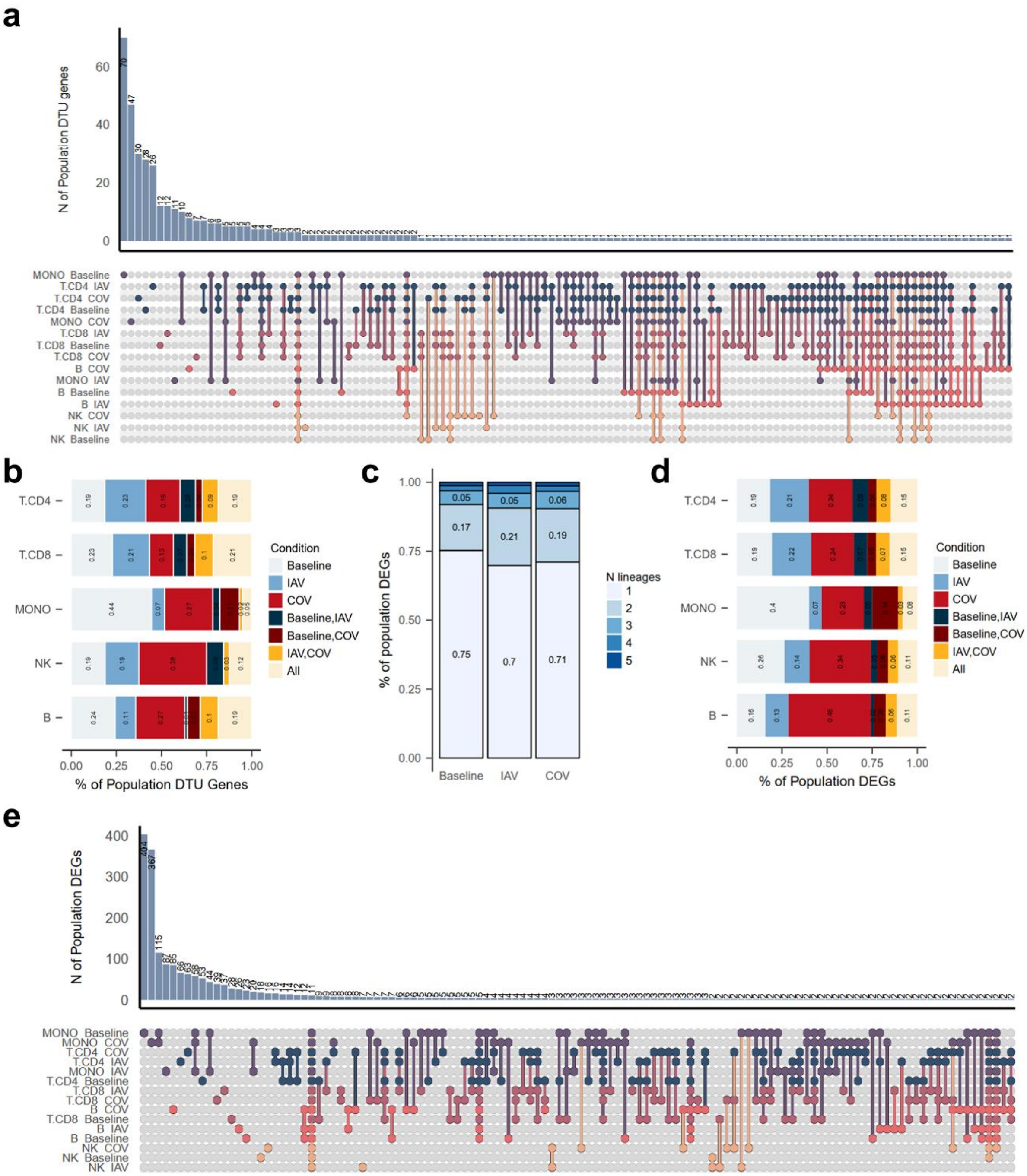


[Supplementary Fig. 10](#). **Sharing of population-level differential transcript usage (DTU) genes and differentially expressed genes (DEGs) across immune lineages and conditions. a)** Upset plot showing the overlap of population DTU genes across immune lineages and conditions (Baseline, influenza A virus (IAV), and SARS-CoV-2 (COV)). **b)** Proportion of population DTU genes detected in each condition combination within each immune lineage. **c)** Proportion of population DEGs shared across immune lineages for each condition. **d)** Proportion of population DEGs detected in each condition combination within each immune lineage. **e)** Upset plot showing the overlap of population DEGs across immune lineages and conditions, considering only gene sets containing >= 1 gene. Across all panels, analyses were restricted to the common set of genes tested across all lineage–condition combinations.

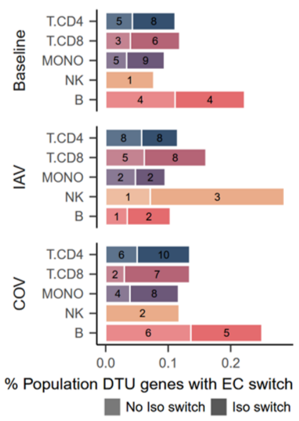


[Supplementary Fig. 11](#). **Proportion of population-level differential transcript usage (DTU) genes that show a switch in the dominant equivalence class (EC) between populations.** Percentage of population DTU genes with a switch in their mean most abundant EC between individuals of European and African populations. EC switches are further stratified between those genes where the switching ECs map to non-overlapping sets of transcripts (Iso swtich; e.g. EC1 maps to t1 and EC2 maps to t2,t3), and those that map to overlapping sets of transcripts (No iso switch; e.g EC1 maps to t1 and EC2 maps to t1,t2).

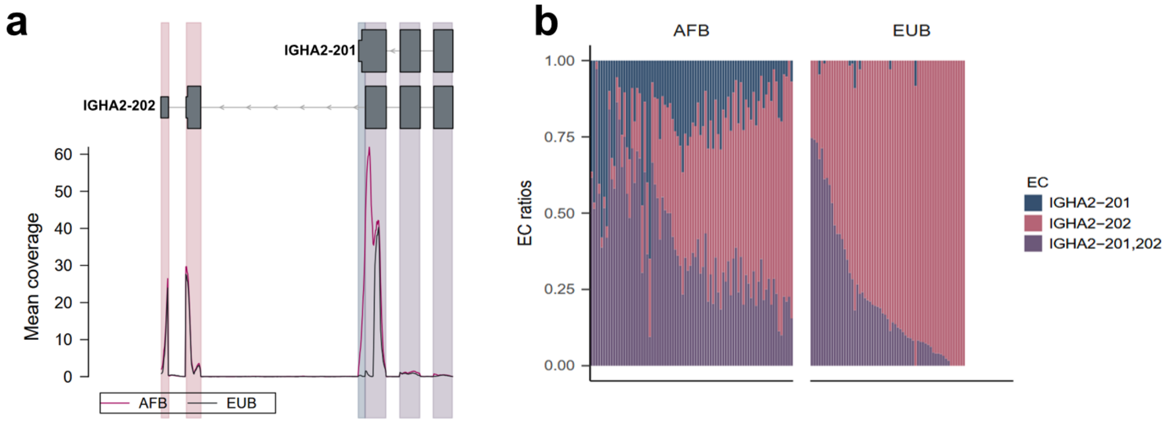


[Supplementary Fig. 12](#). **Population-specific equivalence class (EC) switch at the *IGHA2* locus in resting B cells. a)** Mapping of IGHA2 ECs to transcript genomic coordinates. The three equivalence classes (ECs) correspond to transcript fragments derived from two annotated transcripts (*IGHA2-201* and *IGHA2-202*), shown in grey with UTRs as thin lines and coding sequences as thick lines. Colored vertical blocks indicate the transcript segments represented by each EC *(IGHA2-201*; *IGHA2-202*; *IGHA2-201,202*). Mean coverage profiles across African (AFB) and European (EUB) samples in resting B cells are shown below (10-bp bins). **b)** Relative EC abundances per donor stratified by population. There is a shift in the dominant from *IGHA2-202* in Europeans (mean EUB=0.77; mean AFB=0.39) to IGHA2-201,202 in Africans (mean EUB=0.22; mean AFB=0.40), due to African-specific expression of the IGHA2-201 EC (mean EUB= 0; mean AFB= 0.21).

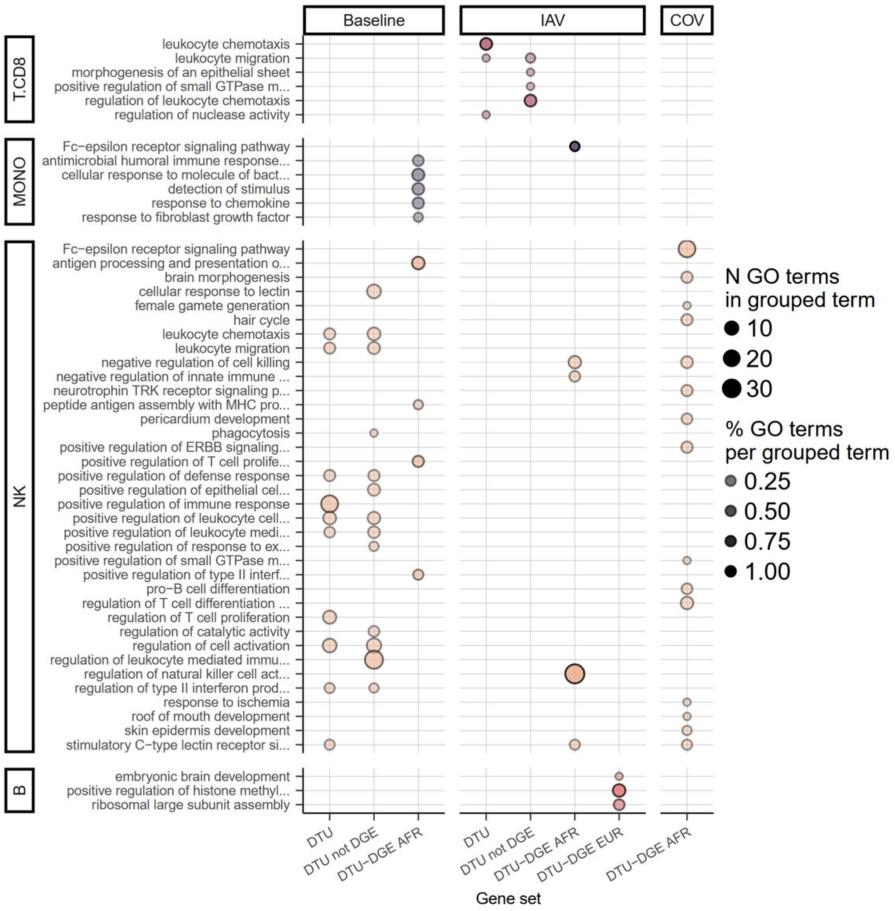


**Supplementary Fig. 13. Pathway overrepresentation analysis (ORA) analysis of population-associated differential transcript usage (DTU) genes.** ORA of population DTU genes identified across immune lineages under baseline, influenza A virus (IAV), and SARS-CoV-2 (COV) conditions. Gene sets include: (i) all DTU genes (DTU); (ii) DTU genes overlapping with differentially expressed genes (DTU & DGE); and (iii) DTU & DGE genes further stratified as upregulated in Africans (DTU & DGE AFR) or upregulated in Europeans (DTU & DGE EUR). Gene Ontology (GO) biological process terms with FDR < 0.05 (Benjamini–Hochberg correction) are shown. GO terms were grouped using rrvgo[4] under a similarity threshold of 0.7. Dot color intensity and size represent the fraction and absolute number of GO terms within each grouped category, respectively.

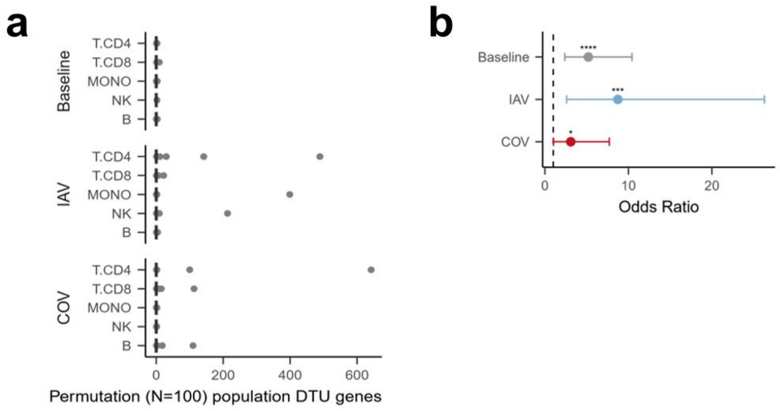


**Supplementary Fig. 14. Validation of population-level differential transcript usage (DTU) results by permutation and comparison with previous studies. a)** Permutation analysis of population DTU. For each immune lineage and condition (e.g., CD4+ T baseline cells), donor labels were randomly shuffled and the DTU analysis was repeated 100 times. Boxplots show the number of significant population DTU genes detected under permutation (FDR < 0.05, Benjamini–Hochberg correction); grey points indicate outliers. **b)** Overlap between monocyte population DTU genes identified in this study and those reported in a bulk RNA-seq study of monocytes[4]. Asterisks denote significance levels (* p < 0.01, ** p < 0.001, etc.).

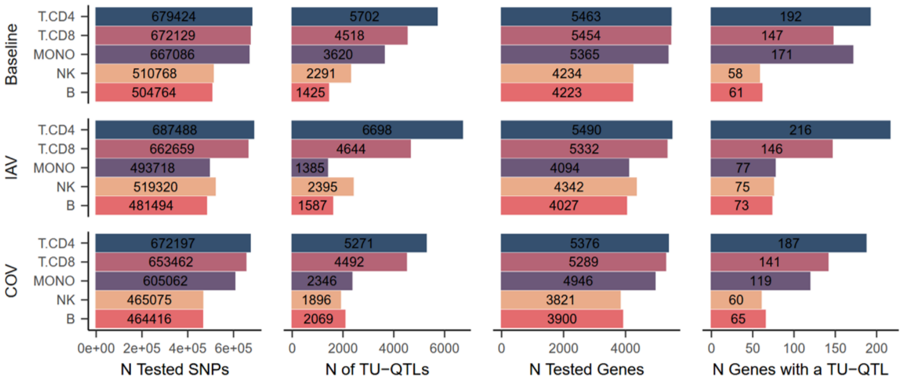


**Supplementary Fig. 15. Summary of tested genetic variants and genes in transcript usage quantitative trait loci (TU-QTL) mapping analysis across immune lineages and conditions.** From left to right, number of tested genetic variants (single nucleotide polimorphisms; SNP), number of significant TU-QTLs (FDR<0.01), number of tested genes, and number of genes with a significant TU-QTL across immune lineages and conditions.

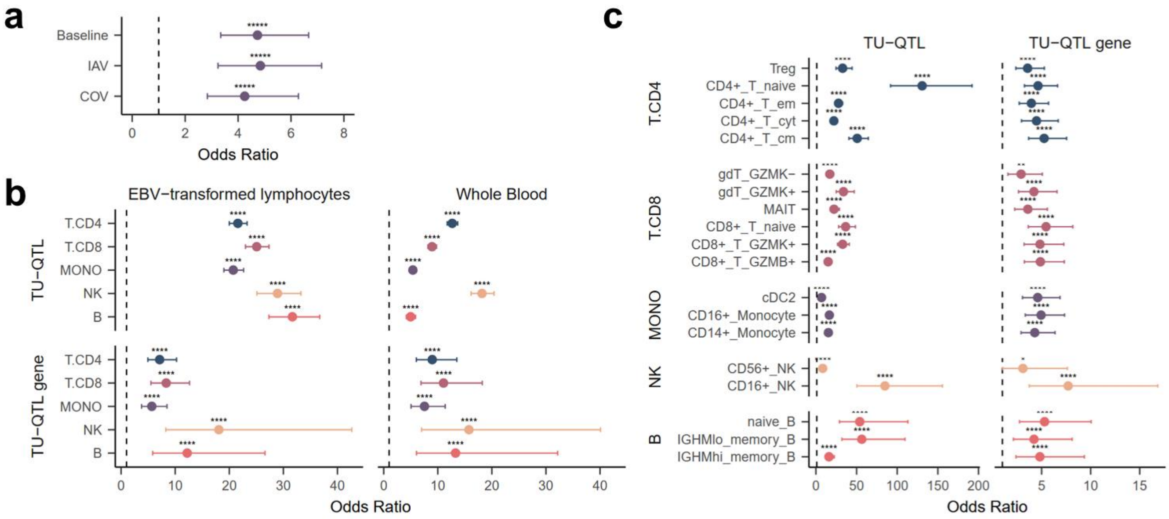


**Supplementary Fig. 16. Overlap of transcript usage quantitative trait loci (TU-QTLs) and associated genes found in our study those detected in previous studies. a)** Overlap between monocyte TU-QTL genes identified in this study and those reported in a bulk RNA-seq study of monocytes[5]. Baseline TU-QTL genes were compared with baseline genes, while influenza A (IAV) and SARS-CoV-2 (COV)-associated TU-QTL genes were compared with IAV-associated genes. The dotted line indicates no enrichment (odds ratio = 1). Asterisks denote significance levels: * p < 0.01, ** p < 0.001, etc (Fisher's exact test). **b)** Overlap between TU-QTLs and TU-QTL genes detected in this study and those reported in bulk RNA-seq whole blood and EBV-transformed lymphocytes[6] **c)** Overlap between baseline TU-QTL genes from this study and those detected in a 5' single-cell study of TU-QTL across Asian populations in immune cells at baseline[7].

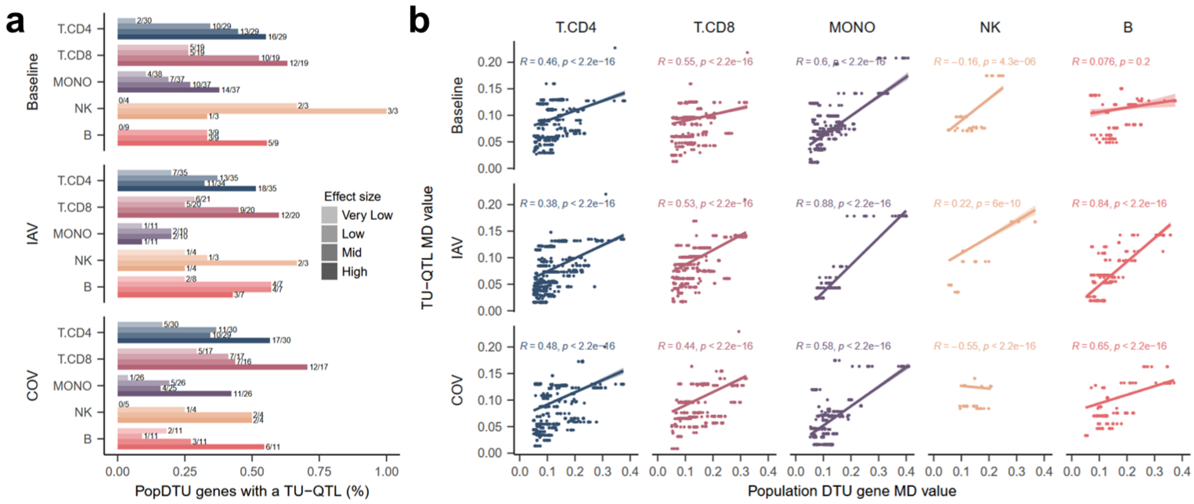


[Supplementary Fig. 17](#). **Population-associated differential transcript usage (DTU) genes with the largest ancestry differences are more likely to be under genetic regulation and associated with large-effect transcript usage quantitative trait loci (TU-QTL). a)** Proportion of population DTU genes harboring a TU-QTL, stratified by TU effect size (maximum absolute difference in mean EC ratio across sample groups, MD value) quartiles, showing enrichment of TU-QTLs among genes with larger ancestry-associated TU differences. **b)** Correlation between TU-QTL effect size (MD value) and DTU effect size (MD value) for population DTU genes with at least one TU-QTL, across immune lineages and conditions. Spearman correlation coefficients (R) and corresponding *p* values are shown.

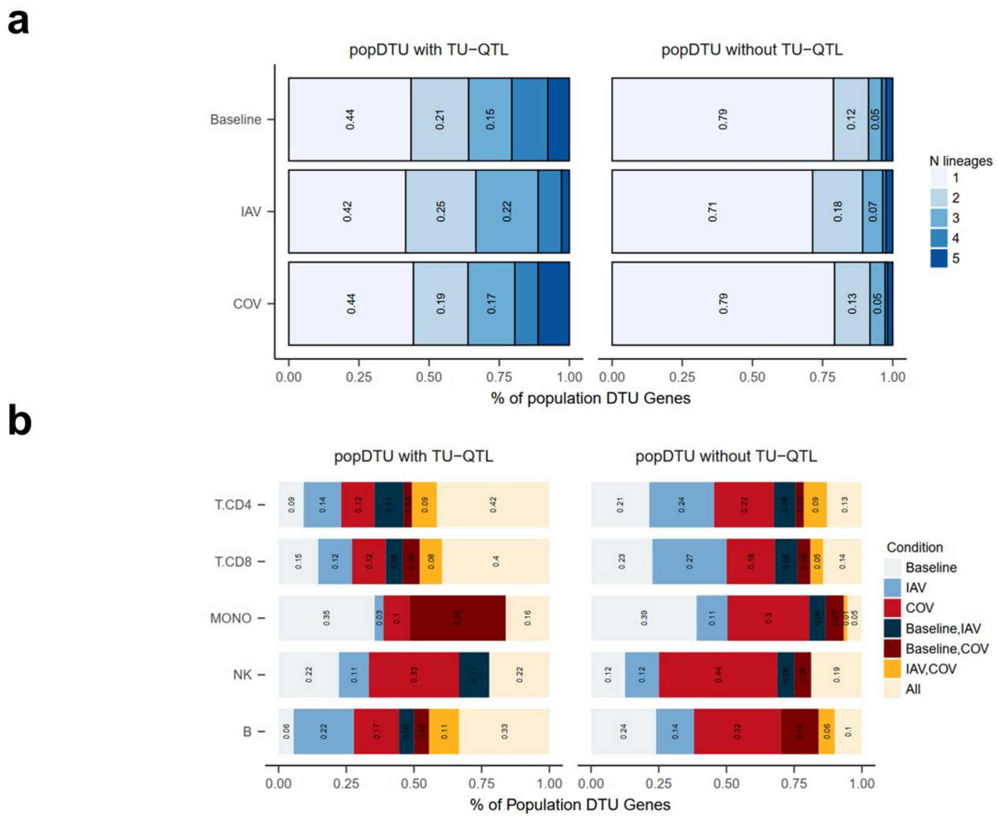


[Supplementary Fig. 18.](#) **Sharing of population differential transcript usage (DTU) genes with and without a transcript usage quantitative trait loci (TU-QTL) across immune lineages and conditions. a)** Proportion of population DTU genes with and without a TU-QTL shared across immune lineages within each condition. **b)** Proportion of population DTU genes with and without a TU-QTL detected in each condition combination within each immune lineage. A common set of tested population DTU genes across immune lineages and conditions is considered across the different plots.

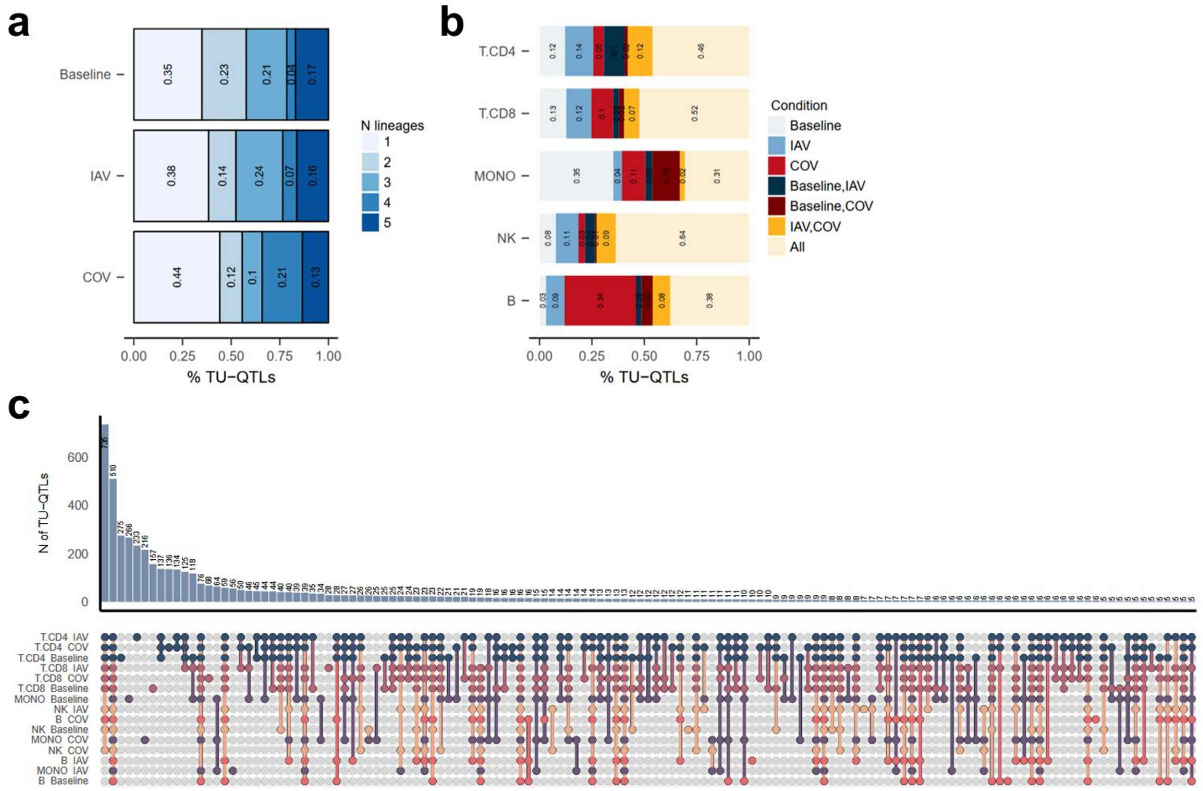


[Supplementary Fig. 19.](#) **Sharing of transcript usage quantitative trait loci (TU-QTLs) across immune lineages and conditions. a)** Proportion of TU-QTLs shared across immune lineages for each condition. **b)** Proportion of TU-QTLs detected in each condition combination within each immune lineage. **c)** UpSet plot showing the overlap of TU-QTLs across immune lineages and conditions for sets >=6 genes. A common set of TU-QTLs tested across immune lineages and conditions is considered across the different plots

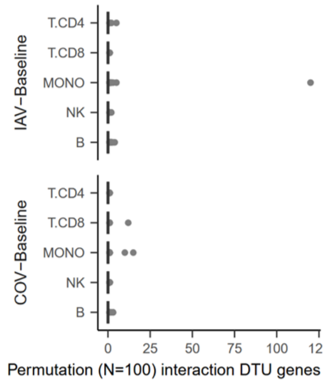


[Supplementary Fig. 21](#)**. Validation of infection x population differential transcript usage (DTU) genes using permutation analysis.** Under the original (non-permuted) infection x population DTU analysis, in each viral response (e.g. influenza A virus (IAV)-Baseline, SARS-CoV-2 (COV)-Baseline), equivalence class (EC) differences were computed between matched infected and baseline samples from the same donor. (e.g. $Donor_{1\ Infected}$-$Donor_{1\ Baseline}$, $Donor_{2\ Infected}$-$Donor_{2\ Baseline}$). For the permutation analysis, instead, we randomly shuffled the condition labels across donors, such that EC ratio differences were calculated between mismatched infected and baseline samples (e.g., $Donor_{1\ Infected}$-$Donor_{1\ Baseline}$, $Donor_{2\ Baseline}$-$Donor_{2\ Infected}$). This procedure was repeated 100 times, and the infection x population DTU analysis was performed for each permutation. Boxplots show the number of significant infection × population DTU genes detected per permutation (FDR < 0.05, Benjamini–Hochberg correction). Center line: median; box limits: upper and lower quartiles; whiskers: 1.5× interquartile range; points: outliers.

**a**

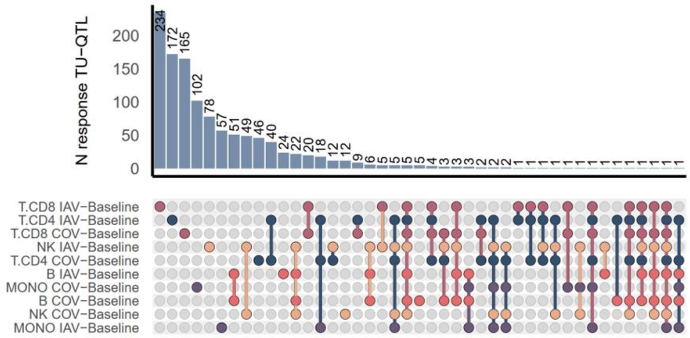


**b**

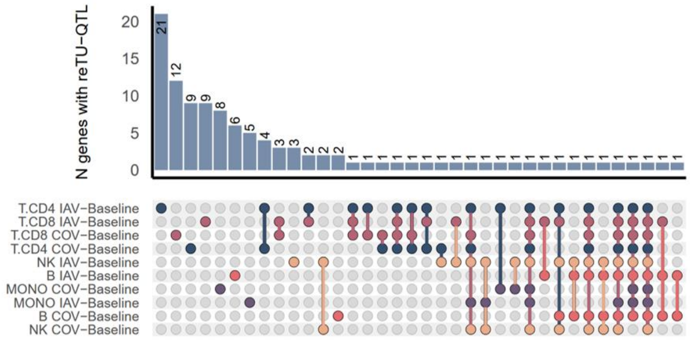


Supplementary Fig. 21. **Sharing of response transcript usage quantitative trait loci (response TU-QTL) across immune lineages and viral stimulations. a)** UpSet plot depicting the sharing of response TU-QTLs across immune lineages and influenza A (IAV-Baseline) or SARS-CoV-2 (COV-Baseline) responses. Response TU-QTLs are defined as SNPs significantly associated with transcript usage variation only after infection ($FDR_{IAV,COV} < 0.01$) but not in baseline ($p_{Baseline} > 0.01$). **b)** UpSet plot depicting the sharing of genes harbouring a response TU-QTL across lineages and viral responses.

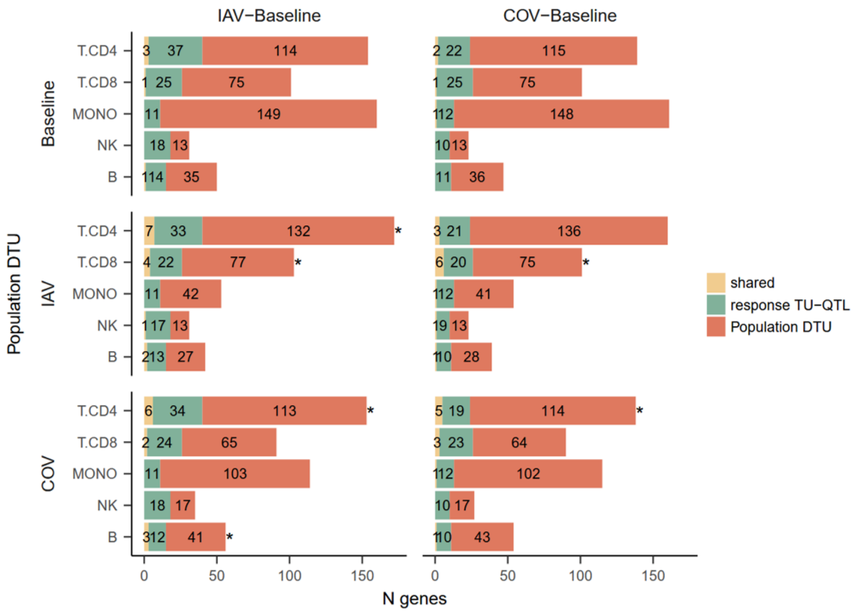


[Supplementary Fig. 22](#). **Overlap of population differential transcript usage (DTU) genes with genes harbouring a response transcript usage quantitative trait loci (re-TU-QTL) across immune lineages and viral responses.** Population DTU genes detected in each separate condition (Baseline, IAV, COV) are overlapped with genes with a response TU-QTL in either influenza A (IAV-Baseline) or SARS-CoV-2 (COV-Baseline) response using a Fisher exact test. Asterisk denotes significant overlaps (FDR< 0.05).

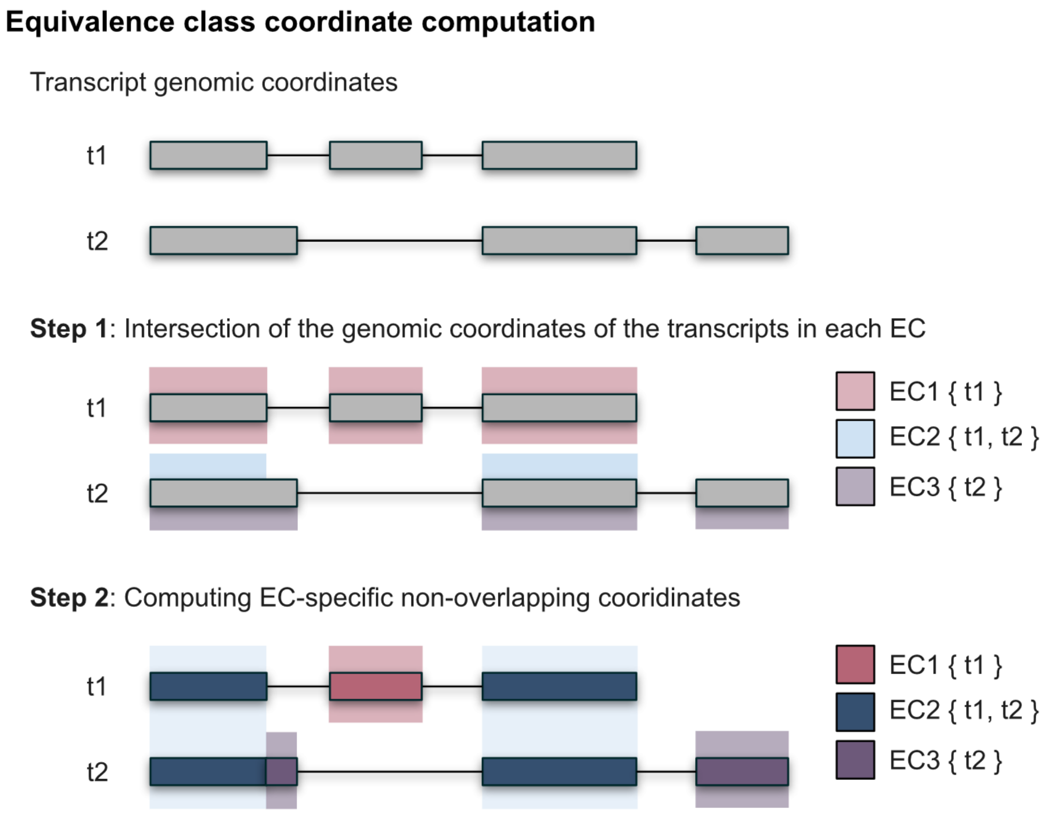


[Supplementary Fig. 23](#). **Computation of genomic coordinates for each equivalence class (EC).** To assign genomic coordinates to ECs, we apply a two-step procedure. Step 1: For each EC, we calculate the intersection of the genomic coordinates of all transcripts it contains. For example, EC1 {t1} is assigned the coordinates of transcript t1; EC2 {t1, t2} is assigned the intersection of t1 and t2 coordinates; and EC3 {t2} is assigned the coordinates of t2. Colors indicate the intersection of the genomic coordinates of the transcripts in each EC. Step 2: To obtain non-overlapping regions specific to each EC, we subtract shared coordinates between ECs. Specifically, the intersected coordinates of EC2 {t1, t2} are subtracted from those of EC1 {t1} and EC3 {t2}, yielding EC-specific non-overlapping genomic segments. Colors indicate the final non-overlapping genomic regions assigned to each EC.
